# Oral Actinobacteria sense and defend against parasitic epibionts

**DOI:** 10.64898/2026.09.08.750141

**Authors:** Yaxi Wang, Jake Colautti, Larry A. Gallagher, Jaehoon Lee, Yongjun Tan, Savannah Farrell, Conor Farrell, Batbileg Bor, John C. Whitney, Dapeng Zhang, S. Brook Peterson, Joseph D. Mougous

## Abstract

Bacteria contend with a wide range of antagonistic interactions from neighboring microbes. One widespread but poorly understood threat is posed by Patescibacteria, which proliferate by colonizing host bacteria surfaces and extracting cellular resources. Despite the ubiquity of Patescibacteria in the environment, how hosts defend against these epibionts remains unknown. Here we show that Actinomycetota fend off Patescibacteria using the Esx secretion system. We identify a secreted effector protein that mediates this defensive behavior and demonstrate that Esx induction is part of a multifaceted, epibiont-specific response that includes multiple predicted cell surface modifications. Finally, we show that this response occurs at the transcriptional level and is mediated by a previously undescribed threat sensing pathway whose activation leads to threonine phosphorylation of an FHA domain-containing output protein. Together with bioinformatic evidence linking this threat sensing pathway to Esx across Actinomycetales, our experimental findings establish a functional role for the elusive and broadly distributed Esx pathway in defense against epibionts. Furthermore, they help resolve the ecological relationship between Patescibacteria and their hosts as one of antagonism, not mutualism.

## Introduction

Immune responses to biological threats, once thought to be confined to eukaryotic organisms, are increasingly recognized as important determinants of bacterial fitness^1,2^. For instance, the burden imposed by phage infection has selected for the expansion and diversification of corresponding defense mechanisms^3^. These pathways are typically discrete functional modules that include a sensor for detecting the presence of an infection coupled to production of one or more proteins that acts to directly inhibit phage proliferation or which causes abortive infection by inhibiting the growth of the infected cell^4^. In contrast, defense against bacterial antagonists can encompass genome wide responses that include induction of both protective mechanisms and counteroffensive antagonistic pathways^2^. *Pseudomonas* species, for example, employ a global posttranscriptional regulatory pathway that alters expression of more than 200 proteins in response to lysis of neighboring kin cells^5–7^. These include pathways that confer protection against specific antibacterial toxins, and the type VI secretion system (T6SS) that can intoxicate competitors^8^. Other antibacterial defense mechanisms bacteria employ include production of protective polysaccharide capsules or extracellular amyloid proteins, induction of stress responses, and deployment of immunity proteins that neutralize incoming toxins and are encoded in actively acquired gene clusters^9–14^.

Patescibacteria are a diverse phylum of ubiquitous but poorly characterized small bacteria with reduced genomes coding for limited anabolic capability^15,16^. A breakthrough in the study of these elusive organisms came with the discovery that Patescibacteria belonging to the Saccharibacteria class, which are present in the oral cavity of >90% of humans, grow epibiotically on host Acinobacterial cells^17^. Subsequent fluorescent and electron microscopy studies suggest this form of growth may be conserved across the Patescibacteria phylum^18–21^. However, many questions remain open regarding the interactions between Patescibacteria and their hosts. For instance, the mechanisms underlying host resource co-option remain undescribed, as do the defensive responses such mechanisms would presumably elicit. Indeed, the fundamental ecological nature of the interaction of Patescbacteria and their hosts has yet to be clarified. The Patescibacteria requirement to extract nutrients from their hosts would suggest they impose a metabolic burden, yet one study found that ammonia secretion by Saccharibacteria protects their host from acid stress encountered in the oral cavity, and another reported that Saccharibacteria infection promotes phage resistance^22,23^. Here, we show that Actinobacteria encode a threat sensing system that induces global transcriptional remodeling in response to Patescibacteria infection. We further show this response includes induction of an Esx system that functions to limit Patescibacteria proliferation. Together, our findings provide compelling evidence that the relationship between Patescibacteria and their hosts is fundamentally parasitic and driving the evolution of tailored host defensive strategies.

## Results

### Saccharibacteria infection induces cell surface remodeling in *Schaalia odontolytica*

Several lines of evidence suggest that Saccharibacteria exhibit a parasitic relationship with their host Actinobacteria^16^. We reasoned that hosts might thereby elaborate defensive measures upon encountering Saccharibacteria. To investigate this, we compared the transcriptome of *Schaalia odontolytica* F0309 (formerly *Actinomyces odontolyticus,* So F0309 hereafter) in the presence and absence of *Nanosynbacter* sp. TM7-008 (TM7-008), a Saccharibacteria able to propagate on its surface (Supplemental Figure 1A, B)^24^. We employed a high multiplicity of infection (MOI) and a short infection period, such that a high proportion of hosts would contact a Saccharibacterial cell and to maximize the probability of observing an initial response rather than a secondary physiological consequence of infection, respectively.

Across three replicates, we observed at least 158 transcripts whose abundance was significantly affected by TM7-008 infection (Figure 1A, Supplemental Figure 2A). The majority of these (131) corresponded to genes with increased expression in the presence of TM7-008. Interestingly, among those genes induced by TM7-008 were six discrete gene clusters with predicted functions strongly associated with the cell envelope (Supplemental Table 1). These include cell wall-associated carbohydrate biosynthesis (OBNOAD_01616-01627, ave log2 fold change 1.7), a group of hypervariable, surface-anchored proteins (OBNOAD_00248-00251, ave log2 fold change 1.7), the Esx secretion system (OBNOAD_00322-00326 and OBNOAD_00329-00338, ave log2 fold change 1.2) and predicted Esx substrates encoded in four distinct gene clusters (OBNOAD_00093-00095, ave log2 fold change 1.2; OBNOAD_00540- 00546, ave log2 fold change 1.5; OBNOAD_01965-01966, ave log2 fold change 1.4; OBNOAD_01994-02007, ave log2 fold change 2.0). Functions pertinent to the cell envelope also extended to genes repressed in the presence of TM7-008, including two operons encoding predicted sortase-anchored fimbrial structures and a predicted adhesin.

**Figure 1.**
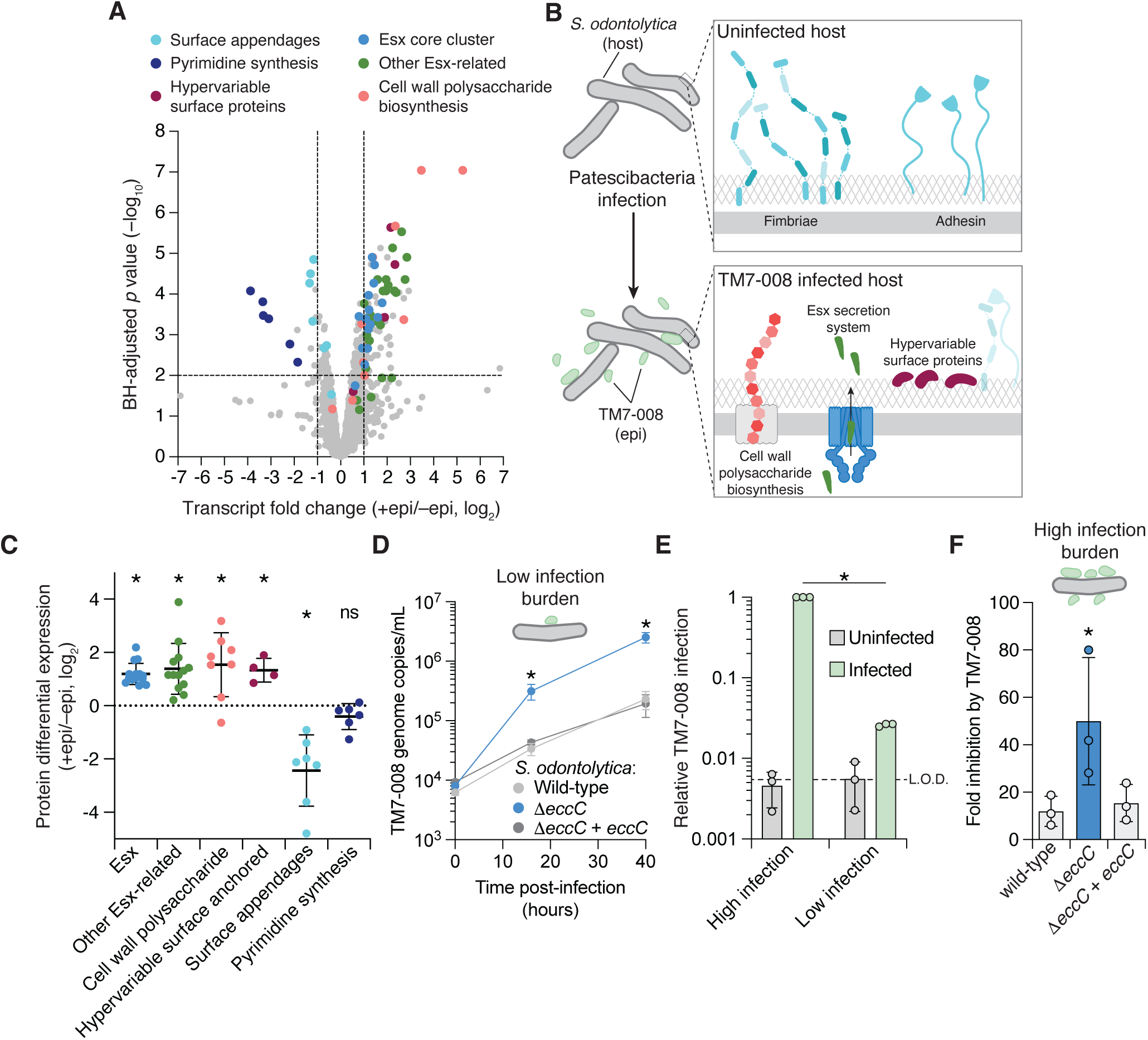
The Esx secretion system of *S. odontolytica* is induced by epibiont infection and inhibits epibiont proliferation. A) Analysis of So F0309 transcript abundance in co-cultures with TM7-008 (+epi) compared to So F0309 mono-cultures (–epi). Gene clusters or groups of functionally-related gene clusters in which multiple genes are significantly affected by TM7-008 infection are highlighted. n = 3 biological replicates. B) Schematic depicting alterations to the So F0309 cell surface expected to result from the transcriptomic changes induced by infection with TM7-008. Coloring scheme is the same as in (A). The fimbrae and adhesin depicted are categorized as surface appendages in (A). C) Differential expression of So F0309 proteins between So F0309–TM7-008 co-cultures and So F0309 mono-cultures. Label-free quantification (LFQ) intensity is normalized by summed LFQ intensity of all So F0309 protein groups in each sample. Missing values in normalized LFQ intensity were imputed as described in Methods. Proteins are grouped and colored in the same way as in (A) and (B). Data represent the mean ± s.d, n = 2 biological replicates. *p≤0.05; ns, not significant, one sample t test comparing mean with value 0. D) Growth of TM7-008 on the indicated *S. odontolytica* F0309 strains, under low infection burden conditions as described in the text. Data represent mean ± s.d. (n = 3 biological replicates). **p≤0.05; two-way repeated-measures ANOVA comparing means at each timepoint. E) Flow cytometric quantification of relative TM7-008 *gfp* infection levels on So F0309 under either high or low infection burden experimental conditions established as described in the text. The dashed line represents the background GFP signal in uninfected control cultures. Data are normalized to the infection level observed under the high infection burden condition and represent mean ± s.d. (n = 3 biological replicates). *p≤0.05; unpaired Welch’s t-test comparing mean relative infection between the high and low burden of infection conditions. F) Growth inhibition (fold change in growth yield) of the indicated So F0309 strains upon infection by TM7-008 under high infection burden conditions, as shown in (E). Data represent mean ± s.d. (n = 3 biological replicates). *p≤0.05; one-way ANOVA with Fisher’s LSD test comparing each mean with the mean of the wild-type control.

### The Esx secretion system of *S. odontolytica* limits Saccharibacteria proliferation

One group of genes activated by TM7-008 that caught our attention were those predicted to participate in the Esx secretion pathway. This system, also referred to as the type VII secretion system, is widespread in Actinomycetota and Bacillota^25^. It consists of the conserved AAA - ATPase (EccC or EssC) required for energizing substrate translocation, one or more homologs of the secreted substrate EsxA, and a membrane pore complex comprised of different proteins in Actinomycetota and Bacillota. In mycobacteria, Esx secretion systems are best known for their role in phagosome escape, nutrient acquisition and conjugative DNA transfer^26,27^. In contrast, the Esx secretion system of Firmicutes is implicated in interbacterial antagonism, and a recent study suggests this function may also extend to some rapidly growing mycobacterial species^28–32^. We therefore reasoned that activation of the Esx pathway in So F0309 could represent an adaptive response of So F0309 to the presence of Saccharibacteria. Supporting this notion, we found that the proteomic response to TM7-008, including the increased production of Esx-related proteins, mirrored many of the transcriptional changes observed in the presence of the parasite (Figure 1B, C, Supplemental Figure 2B, Supplemental Table 2).

To probe the functional consequence of Esx in the context of Saccharibacterial–host interaction, we sought to genetically inactivate the pathway. Towards this end, we developed a new counter-selectable allelic exchange plasmid for use in *Schaalia*, *Actinomyces* and related species, pLG202 (Supplemental Figure 3A). This plasmid incorporates a modified phenylalanyl-tRNA synthetase allele with relaxed substrate specificity (*pheS*^T275A, A322G^), enabling counter-selection against the integrated vector with p-chlorophenylalanine (Supplemental Figure 3B). With this tool in hand, we generated an in-frame deletion of *eccC,* encoding the conserved AAA+ family ATPase required for activity of the secretion system, and compared the growth of TM7-008 on this mutant and wild-type So F0309. Strikingly, we observed that TM7-008 replication on So F0309 is markedly enhanced when the Esx secretion system of the host is inactivated (Figure 1D). Genetic complementation of *eccC* returned TM7-008 proliferation to wild-type levels. The phenotype we observed is not explained by host cell availability, as So F0309 wild-type and Δ*eccC* grew equivalently in these experiments (Supplemental Figure 4A). Rather, the data suggest that the Esx secretion system of So F0309 limits TM7-008 proliferation.

If the physiological role of the Esx secretion system in *S. odontolytica* is providing defense against Saccharibacteria, we would predict that its inactivation should result in reduced fitness under conditions where Saccharibacterial infection impacts host proliferation. To assess this, we developed a culturing regimen in which TM7-008 imposes a high infection burden upon So F0309. At this higher level of infection (∼40-fold beyond that where TM7-008 proliferation is measured), we found that the growth yield of So F0309 was suppressed ∼10-fold compared to that of uninfected cultures (Figure 1E, Supplemental Figure 4B, C). Under these conditions, inactivation of the Esx secretion system resulted in a ∼50-fold reduction of So F0309 growth yield, consistent with the pathway contributing to defense against the epibiont. (Figure 1F)

### An Esx-secreted effector mediates the effect of the Esx pathway on epibiont proliferation

Next we sought to identify Esx substrates that could mediate its effects on TM7-008. Notably, Esx effectors of a bacterium belonging to the family Actinomycetaceae, which includes the genera *Actinomyces* and *Schaalia*, have not been identified experimentally. Esx secretion system gene clusters typically encode one or more small helical substrates belonging to the WXG100 family, and two of these sharing sequence homology with EsxA (ESAT-6) and EsxB (CFP-10) of *Mycobacterium tuberculosis* were readily identifiable in So F0309 (OBNOAD_00329-00330). However, these proteins typically lack clear effector function and are required for secretion of other substrates^33,34^. Supporting a role for WXG100 family proteins of So F0309 in the basal activity of the apparatus, AlphaFold modeling indicates they form a high-confidence heterodimer with strong structural similarity to the EsxA–EsxB heterodimer required for substrate secretion by ESX-1 of *M. tuberculosis* (RMSD 1.183 Å for 73 pruned amino acid pairs, Supplemental Figure 5A). On the basis of this similarity and shared sequence identity with their *M. tuberculosis* homologs, we have renamed these proteins EsxA and EsxB. Outside of these proteins, So F0309 lacks sequence level homologs of Esx substrates from other species.

A common structural feature shared by Esx substrates from diverse bacteria is the presence of a distinctive N-terminal domain comprised of 110-180 amino acids that adopts an α- helical bundle fold and mediates critical interactions with co-secretion partners^25,35^. Using AlphaFold modeling, we identified four proteins encoded within the So F0309 Esx gene cluster that shared this characteristic (Supplemental Figure 5B). Genome-wide application of this method identified eight additional candidate substrate genes, including five that are transcriptionally activated upon infection by TM7-008. Adding credence to these predictions, each candidate substrate encoded outside of the Esx gene cluster is located adjacent to a gene encoding a small a α-helical harpin protein resembling WXG100 proteins, a genomic arrangement analogous to effectors and their cognate targeting factors, respectively, in the Esx secretion system of Firmicutes^31^.

Given the multiplicity of candidate Esx substrates within So F0309, we sought to experimentally define those secreted under the conditions of our growth assays using an unbiased, semi-quantitative mass spectrometry-based approach. Comparison of the extracellular proteomes of wild-type and Δ*eccC* strains of So F0309 revealed four proteins – all encoded within the Esx gene cluster – whose apparent secretion is diminished by Esx inactivation (Figure 2A). These include EsxA, EsxB and two proteins our structural informatics analyses had identified as candidate substrates (OBNOAD_00323 and 00324, Supplemental Figure 5B). Given the role of EsxA and EsxB in basal Esx function, we focused subsequent phenotypic studies on OBNOAD_00323 and 00324, which we renamed *aesA* and *aesB*, respectively. Inactivation of either candidate substrate compromised the ability of So F0309 to interfere with TM7-008 proliferation (Figure 2B). This phenotype could be genetically complemented and we further found that inactivation of a candidate substrate not detected in our secretomic studies elicited no impact on TM7-008 proliferation (Supplemental Figure 6A).

**Figure 2.**
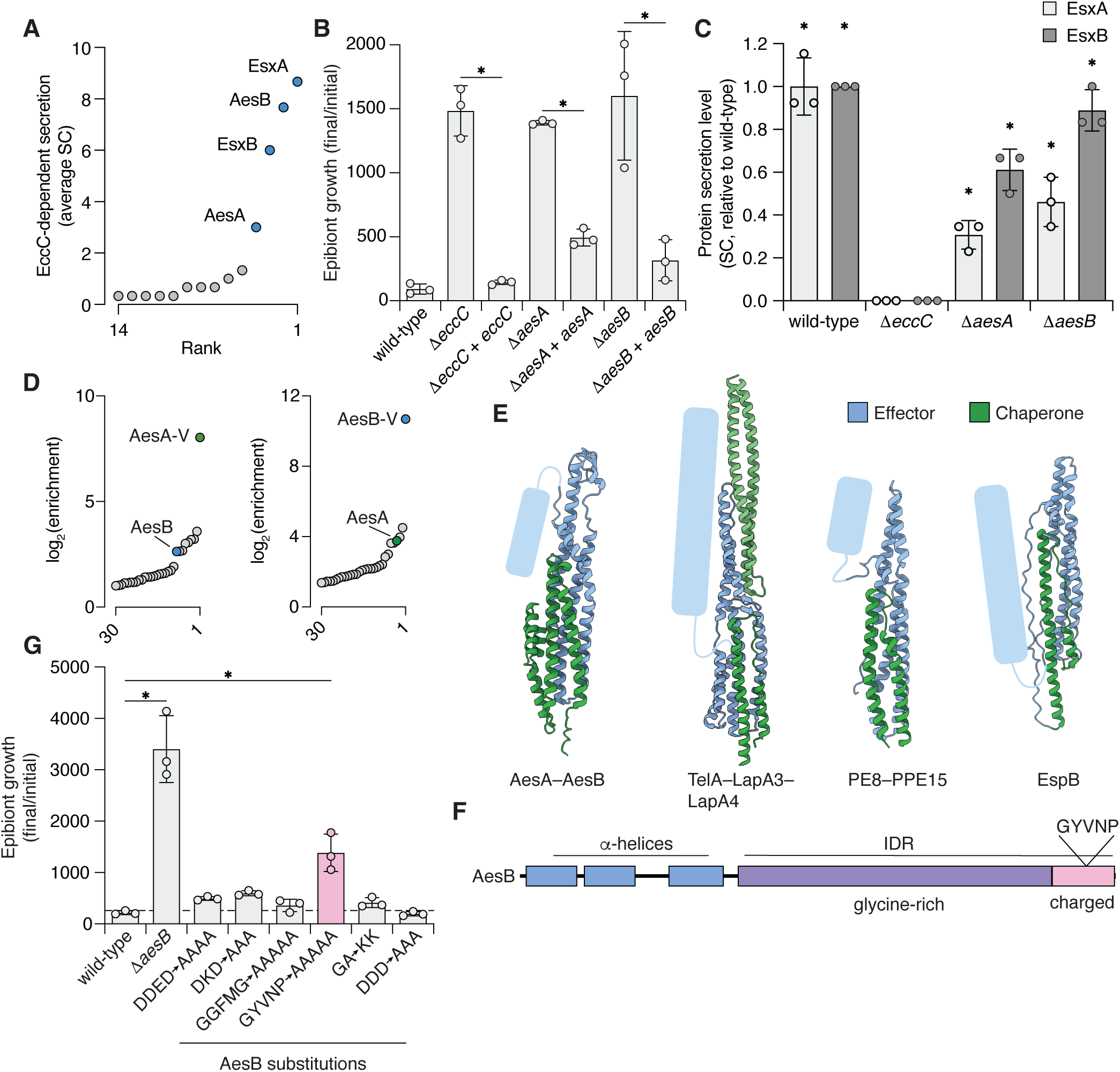
Esx-secreted protein AesB mediates the inhibition of Patescibacteria proliferation. A) Abundance, ranked by mean number of spectral counts detected in mass spectrometry-based analysis, of proteins detected in spent culture supernatant of wild-type So F0309 and missing from So Δ*eccC*. n = 3 technical replicates. B) Change in abundance of TM7-008 after 40 hours of growth on the indicated So F0309 strains. Data represent mean ± s.d. (n = 3 biological replicates). *p≤0.05, one-way ANOVA with Šidák correction for multiple comparisons between the mean of each deletion and the mean of its corresponding complementation strain. C) Level of EsxA and EsxB detected in cell-free culture supernatant of the indicated So F0309 strains, divided by the average spectral counts for each protein in wild-type culture supernatant. Data represent the mean ± s.d. One-way ANOVA with Šidák correction for multiple comparisons between each mean and the mean of the corresponding protein in Δ*eccC*; *p≤0.05 (n = 3 technical replicates). Data in (A) and (C) were obtained from the same experiment. D) Immunoprecipitation-tandem mass spectrometry (IP-MS) identification of proteins that interact with AesA (left) and AesB (right) from So F0309. For each protein, average fold enrichment of proteins in the experimental over the control condition are shown. The enrichment of proteins undetected in control was determined by imputation of the limit of detection for these proteins. Proteins unrelated to the Esx system are shown in grey. n=2 technical replicates. V, VSV-G. E) AlphaFold3 model of the complex predicted to form between AesA and the N-terminal domain of AesB (ipTM = 0.70) compared to characterized complexes formed by other Esx substrates and their co-secreted chaperones. The TelA-LapA3-LapA4 (PDB 8GMH) complex from *Streptococcus intermedius* and PE8-PPE15 (PDB 5XFS) and EspB (PDB 4WJ1) substrates from *M. tuberculosis* are shown for comparison. The unresolved or predicted disordered C-terminal domains of these proteins are schematized as blue oblong shapes. F) Schematic representation of the domain architecture of AesB. A conserved motif in the C-terminal region that is required for function (GYVNP) is highlighted. IDR, intrinsically disordered region. G) Fold change in abundance of TM7-008 after 40 hours of infection on So F0309 strains containing the indicated substitutions in AesB, compared to the wild-type and Δ*aesB* strains. Error bars represent mean ± s.d. (n = 3 biological replicates). *p≤0.05, one-way ANOVA with Šidák correction for multiple comparisons between each mean and the mean of the wild-type control.

The consequences of *aesA* and *aesB* deletion on TM7-008–So 0309 growth dynamics could result from a role of the proteins in directly mediating impacts on TM7-008. Alternatively, AesA and AesB may be required for secretion of other substrates, a phenomenon observed for many of the Esx substrates of *Mycobacterial* species^25^. To distinguish between these possibilities, we evaluated the impact of AesA and AesB inactivation on the secretion of other Esx substrates. Mass spectrometry analysis of culture supernatant derived from So F0309 Δ*aesA* and Δ*aesB* revealed that EsxA and B remain readily detected in these backgrounds (Figure 2C). On the contrary, AesA was not detectable in culture supernatant derived from So F0309 Δ*aesB*, and AesB was undetectable in supernatant from So F0309 Δ*aesA* (Supplemental Figure 6B). In total, these data are consistent with AesA and AesB functioning in an inter-dependent manner downstream of the basal activity of the secretory machinery, conceivably as effectors.

The secretion co-dependence we observed for AesA and AesB is reminiscent of Esx substrates in other organisms. In most cases, this is explained by heterodimerization facilitating secretion^25^. We observed AesB as the one of the most enriched proteins to immunoprecipitate with AesA from So F0309 cellular lysate, and AesA was highly enriched among proteins immunoprecipitating with AesB (Figure 2D). Additionally, AlphaFold predicts a high confidence model in which the N-terminal helical domains of the two substrates interact to form an extended α-helical bundle (Figure 2E, Supplemental Figure 5B). This structure bears notable similarity to complexes formed by characterized Esx secretion substrates, including PE/PPE protein heterodimers and the single protein substrate EspB from *Mycobacterium* species and an Esx substrate in complex with secretion targeting factors LapA3 and LapA4 from *Streptococcus intermedius*^35–40^ (Figure 2E).

In our structural comparisons of the AesA–AesB heterodimer with other Esx substrate pairs, the C-terminal region of AesB analogous to the predicted or established effector domain of other Esx substrates could not be predicted by Alphafold. This portion of AesB contains a ∼160 amino acid region highly enriched in glycine residues, followed by a 56 residue region dominated by charged residues (Figure 2F). Consistent with these observations, algorithms for detecting intrinsically disordered regions (IDRs) predict this portion of the protein is unlikely to adopt a stable conformation (Supplemental Figure 6C). Interestingly, the C-terminal predicted effector or functional domains of many Mycobacterial PPE proteins also contain extended IDRs^41^. Although the function of most IDRs in bacterial proteins remains poorly understood, IDRs are common in eukaryotes, where they often serve as interaction hubs for a variety of macromolecules, including nucleic acids and proteins^42,43^. We thus hypothesized that the C-terminal region of AesB is an important contributor to Esx-mediated inhibition of epibiont proliferation.

Though IDR function typically stems from general properties such as flexibility, charge, charge distribution and hydrophobicity, evolutionary analyses suggest they can also contain regions of sequence conservation linked to their function^44,45^. We used PSI-BLAST and phylogenetic analyses to identify a monophyletic clade of 104 proteins sharing the same domain architecture and a minimum of 30% overall sequence identity with AesB of So F0309 (Supplemental Figure 7A). Multiple sequence alignment of these proteins revealed that while their glycine-rich regions is highly variable in length and sequence composition, their C-terminal charged regions contain multiple conserved sequence motifs (Figure 2F, Supplemental Figure 7B, C). To probe the importance of specific amino acid sequences within these regions, we generated strains expressing AesB variants bearing substitutions in its glycine-rich and charged regions from the native *aesB* locus. We found that non-conservative substitutions in the variable glycine-rich region of AesB or to subsets of charged residues near the C-terminus had no detectable effect on TM7-008 proliferation, a finding in line with the overall character of IDRs being more important than individual residues (Figure 2G). However, a strain expressing AesB with substitutions in a highly conserved motif (GYNVP) permitted significantly more TM7-008 growth than the wild-type, approaching levels of the Δ*aesB* strain. While we were unable to elucidate a mechanism of action for AesB, based on reports wherein small conserved motifs embedded within other IDRs promote docking or complex formation with interaction partners, we speculate that this motif may facilitate binding of a target molecule on TM7-008^46^. Altogether, our results indicate that effector delivery by the Esx secretion system of So F0309 plays an important role in defending against parasitic Saccharibacteria species.

### Identification of a signaling pathway that mediates Esx activation in response to epibiont colonization

We postulated that the transcriptomic and proteomic changes accompanying exposure of So F0309 to TM7-008 could reflect triggering of a specific threat-sensing mechanism. To identify candidates for such a pathway, we searched So F0309 for genes implicated in sensing or responding to biological conflict. We noted that Aravind and colleagues identified predicted threat sensing and response elements composed of, among other proteins, a MoxR-family AAA+ ATPase, a vWA (von Willebrand factor A) domain protein, and variable sensing proteins that can include β-propeller domains^47^. The authors also defined effector functions associated with these elements, including polymorphic toxins and proteins involved in signal transduction such as those containing serine-threonine kinase and forkhead-associated (FHA) domains.

We identified two regions within the So F0309 genome encoding proteins that share hallmarks of the elements identified by Aravind and colleagues (Figure 3A-C)^47^. These include vWA and MoxR proteins, serine-threonine kinases, FHA domain-containing proteins, and predicted cell surface-localized filamentous proteins bearing terminal β-propeller domains.

**Figure 3.**
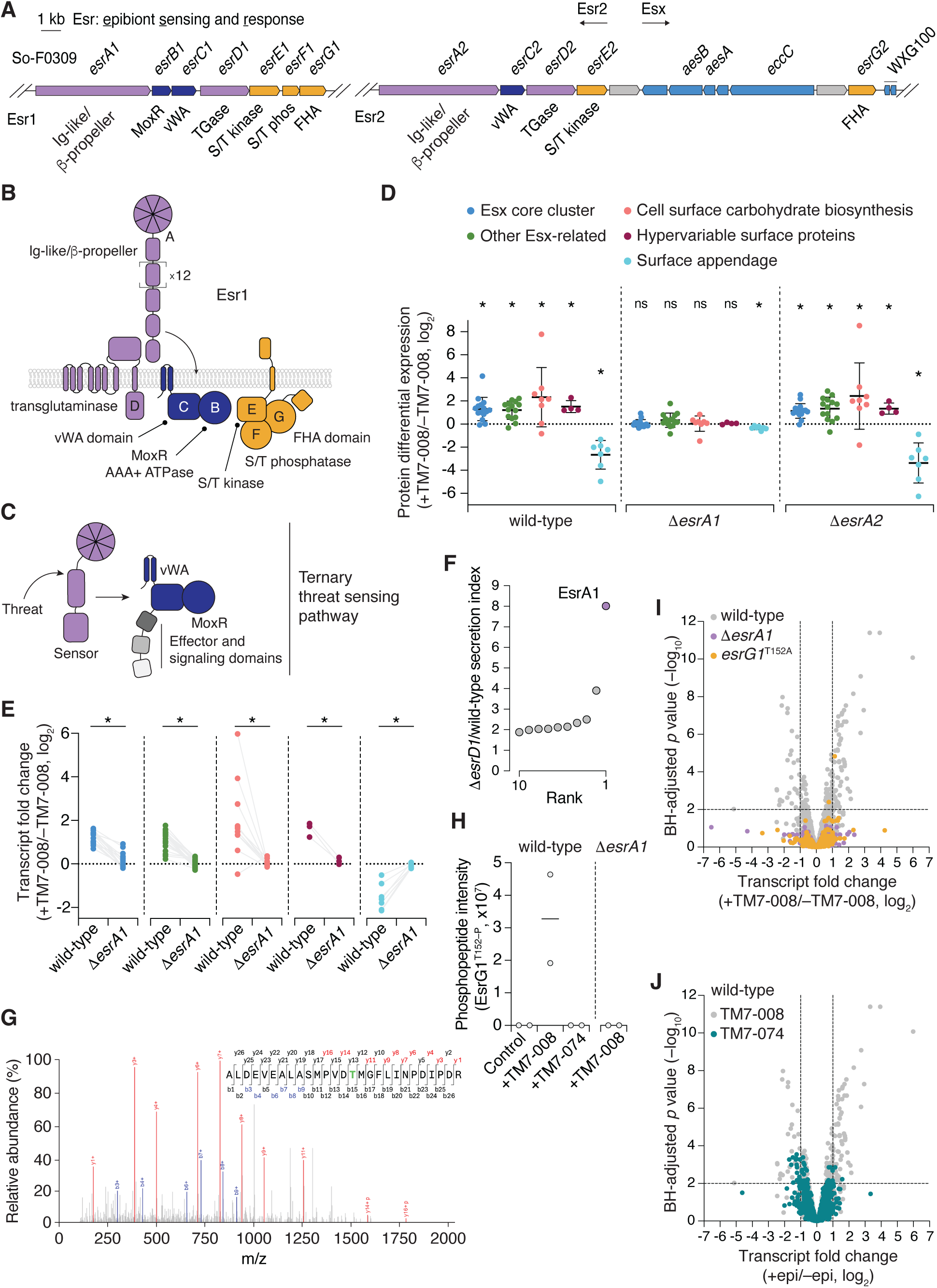
Identification of a threat-sensing pathway in *S. odontolytica* F0309 that is required for Patescibacteria infection-depended responses. A) Schematic depicting the two Esr gene clusters in So F0309. Genes are colored based on functional groups: purple, predicted signal detection and transduction module; yellow, phosphorelay components; blue, Esx substrates and structural elements; grey, other. B, C) Schematic showing predicted protein topology of So F0309 Esr1 components (B) compared to analogous components of ternary threat sensing systems predicted by Aravind and colleagues (C)^47^. Coloring in (B) and (C) is according to (A). S/T, Serine/Threonine. D) Ratio of protein abundance in the indicated strains of So F0309 co-cultured with TM7-008 compared to mono-cultures. Protein levels represent averaged, normalized LFQ values as described in Figure 1C. Data represent the mean ± s.d. for proteins in the indicated category, n = 2 biological replicates. *p≤0.05, one sample t test comparing mean with value 0; ns, not significant. E) Log_2_ fold change in selected So F0309 transcript levels between So F0309–TM7-008 co-cultures and So F0309 mono-cultures in So F0309 wild-type and So F0309 Δ*esrA1.* Lines connect the same transcript between strain backgrounds. n = 3 biological replicates. *p≤0.05, paired two-tailed student t test comparing mean values for all transcripts in the category. F) Rank secretion index plot indicating proteins most differentially abundant (based on MS spectral counts) in cell-free culture supernatant of F0309 Δ*esrD1* compared to wild-type. Data were normalized to spectral counts obtained from cell-associated fractions. G) Tandem mass spectra of the indicated So F0309 peptide from an So F0309 wild- type–TM7-008 co-culture. Matched fragmentation ions (N-terminal b-ions in blue and C- terminal y-ions in red) and the site of phosphorylation (green; T152 in EsrG1) are indicated. H) EsrG1 T152 phosphorylation in So F0309 wild-type or Δ*esrA1* grown in mono-cultures (Control) or co-cultures with TM7-008 or TM7-074, based on MaxQuant phosphor (STY) analysis of whole cell proteome datasets. The line indicates the mean. n = 2 biological replicates. Values are raw MS1 intensities (arbitrary units). I, J) Analysis of So F0309 transcript abundance in co- cultures with TM7-008 (I) or TM7-074 (J) compared to So F0309 mono-cultures. Data obtained from the indicated mutant strains (I) or during co-culture with TM7-074 (J) are overlayed on wild-type ± TM7-008 values. n = 3 biological replicates.

Remarkably, one of these clusters is integrated into the operons encoding the Esx secretion system. Despite their compelling similarity, the gene clusters we identified were not detected in the Aravind study. Notably, the authors of that study initially demarcated genomic regions for in-depth query by the presence of polymorphic toxins and a subset of intermicrobial conflict-associated domains that are not present in the gene clusters we identified. Nevertheless, we hypothesized that the So F0309 gene clusters encode a pathway for sensing and responding to epibiont colonization, and accordingly named their constituents epibiont sensing and response (*esr*) genes.

To test the significance of the Esr pathways during epibiont infection, we compared the proteomic response to TM7-008 colonization of So F0309 wild-type and strains bearing in-frame deletions of each *esrA* gene. Whereas So F0309 lacking *esrA* adjacent to the esx genes (Δ*esrA2*) shared the strong proteomic response of wild-type to TM7-008, the Δ*esrA1* strain displayed virtually no significant proteomic changes associated with TM7-008 infection (Supplemental Figure 8A). Esx proteins, cell wall-associated polysaccharide biosynthesis machinery, hypervariable surface proteins and extracellular appendages were all no longer significantly altered in abundance during TM7-008 infection in So F0309 Δ*esrA1* (Figure 3D). In support of the importance of Esr1 for sensing and responding to TM7-008, we found that deletion of *esrA1* substantially reduced the transcriptomic changes associated with epibiont infection, including eliminating the induction of cell wall-associated polysaccharide biosynthesis and Esx pathway genes (Figure 3E).

The *esrA1* gene encodes a protein with several features suggestive of a role in sensing an extracellular signal, including a predicted sec secretion signal, a series of 14 repeated Ig-like domains predicted to form a filamentous structure, and a N-terminal β-propeller domain from the WD40 class typically involved in protein-protein interactions^48^. Although this protein lacks the LPTXG motif associated with peptidoglycan anchored proteins in monoderm bacteria, we noted that both *esrA* gene clusters encode a predicted transglutaminase protein (EsrD), which we hypothesized could be employed in covalently linking EsrA to a surface component. Supporting this, we found using semiquantitative mass spectrometry that – proteome wide – EsrA1 is the protein most released to the culture supernatant in So F0309 Δ*esrD1* relative to wild-type (Figure 3F).

The presence of genes encoding serine/threonine protein kinases (*esrE1* and *esrE2*), and a phosphatase (*esrF1*) in the esr gene clusters suggested they transduce the sensing of N1-008 infection through protein phosphorylation. To assess this, we examined the proteome of So F0309 for proteins phosphorylated in response to TM7-008. The sole differential phosphorylation signal we detected in these samples was found on threonine 152 of EsrG1 (Figure 3G). This phosphorylation event was abolished upon inactivation of EsrA1, consistent with the sensing of epibiont infection by EsrA1 triggering kinase activation (Figure 3H). EsrG1 is comprised of an N-terminal pseudophosphatase domain (aa1-136), a middle predicted disordered region that includes T152 (aa137-328), and a C-terminal FHA domain (Supplemental Figure 8B). FHA domains typically function by binding phosphothreonine residues^49^, suggesting that phosphorylation of EsrG1 drives a conformational change in the protein that ultimately results in pathway activation. In line with this model, we found that substitution of T152 with alanine largely ablates the transcriptional impact of TM7-008 on So F0309, phenocopying the impact of EsrA1 inactivation (Figure 3I).

Altogether, our findings point to a model in which EsrA1 – covalently linked to an unknown structure on the cell surface by EsrD1 – senses the presence of an infecting epibiont, triggering activation of the kinase EsrE1, which phosphorylates EsrG1, ultimately resulting in transcriptional remodeling that affords protection against epibiont infection. One predicted consequence of So F0309 harboring a pathway that defends against infecting epibionts is selection for epibiont strains that evade Esr-mediated defenses. *Nanosynbacter* sp. TM7-074 is an oral Saccharibacterium that, like TM7-008, replicates epibiotically on So-F0309 (Supplemental Figure 1A). We found that in contrast to TM7-008, TM7-074 replicates equally well on wild-type and Esx-inactivated So F0309 (Supplemental Figure 8C). Additionally, transcriptomic profiling of So F0309 exposed to TM7-074 revealed that, contrary to the robust response to TM7-008 infection, few genes exhibited altered expression levels in response to this species (Figure 3J). This finding suggests that TM7-074 fails to trigger activation of the Esr pathway. Phosphoproteomic analysis demonstrated that indeed, EsrG1 does not become phosphorylated during TM7-074 infection (Figure 3H). TM7-008 and TM7-074 are distinct species, sharing an average nucleotide identity of 82%. Moreover, they encode 145 and 122 proteins with no corresponding orthologs, respectively, precluding straightforward identification of the factor(s) driving these differences. Nonetheless, the evasion of detection by Esr1 display by TM7-074 suggests an arms race dynamic fueling is the evolution of conflict-mediating pathways in Saccharibacteria and their hosts.

### Esr pathways are linked to Esx genes and broadly distributed across Actinomycetes

The range of host species colonized by Patescibacteria remains poorly understood; however, studies suggest that in the oral cavity, diverse Actinobacteria can support the growth of one or more epibiotic species^24,50^. To determine whether the Esr-mediated response to epibiont infection is conserved, we performed whole cell proteomic analyses of a second Actinobacterial species that both encodes the Esr1 pathway and supports replication of TM7-008, *Schaali meyeri* W712 (Supplemental Figure 1A). Our data showed that despite the phylogenetic distance between the strains, TM7-008 infection of *S. meyeri* W712 yields a response, albeit muted, closely resembling that elicited in So-F0309: activation of Esx and cell wall-associated polysaccharide biosynthetic pathways, and repression of orthologous fimbrial structures (Figure 4A, Supplemental Figure 8D). Furthermore, as in TM7-008, these proteins account for a significant fraction of the proteins most differentially expressed in response to the epibiont (10 of 39 proteins more than 2-fold different, compared to 28 out of 70 in So F0309). We found *S. meyeri* intractable to genetic manipulation and were unable to inactivate its Esr pathways; however, phosphoproteomics identified an TM7-008-dependent threonine phosphorylation site (T263) in the EsrG1 protein of this strain (Figure 4B., Supplemental Figure 8E). Notably, this residue is found in the same predicted IDR as the corresponding phosphorylated residue of So F0309 EsrG1. These data support the Esr pathway widely participating in epibiont sensing and mediating a genome-wide response with many conserved elements.

**Figure 4.**
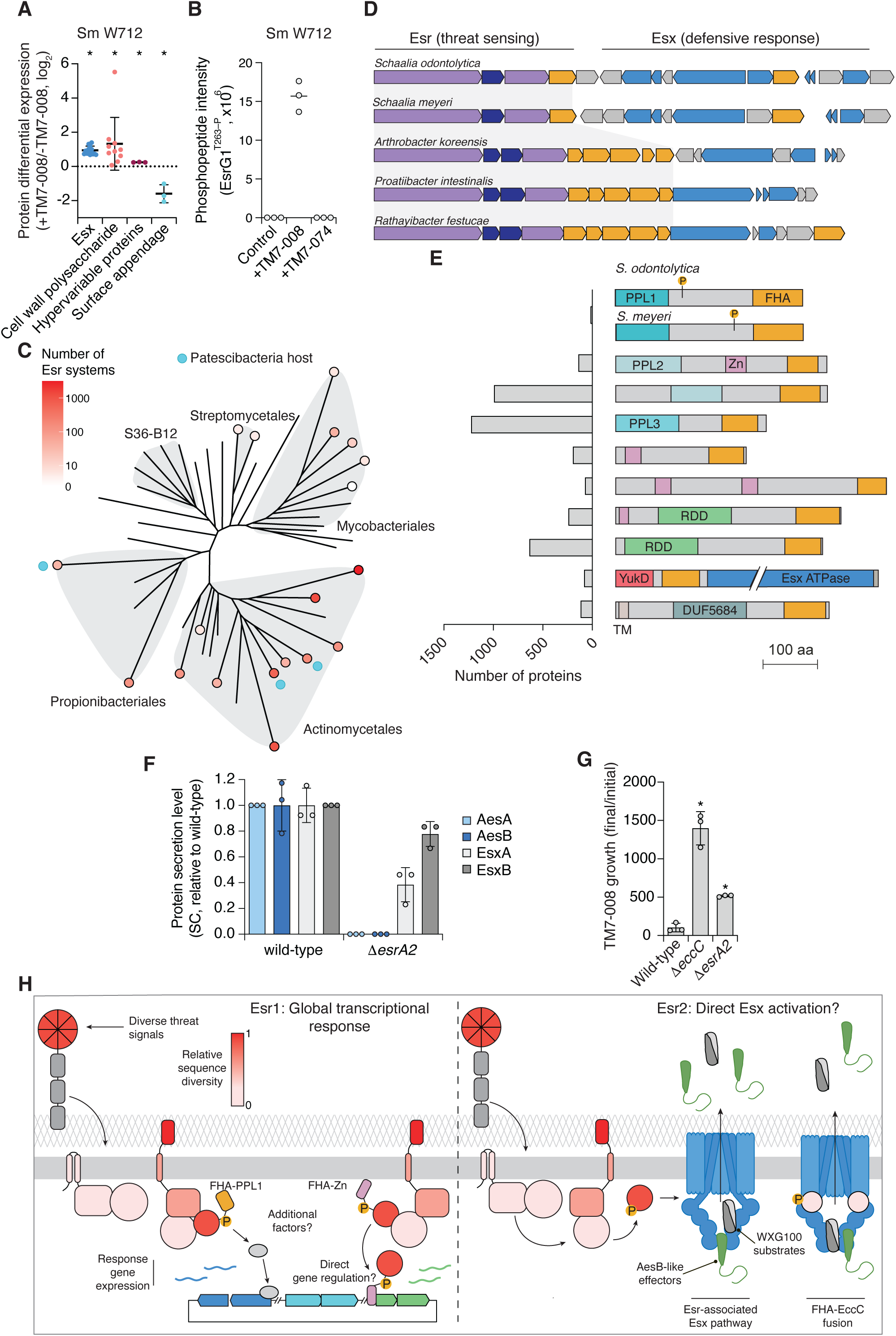
Esr pathways are broadly distributed in Actinomycetes and frequently linked to Esx secretion systems. A) Differential expression of *S. meyeri* W712 proteins between W712– TM7-008 co-cultures and W712 mono-cultures. Protein levels represent averaged, normalized LFQ values as described in Figure 1C. Proteins are grouped and colored in the same way as in Figure 1 and Figure 3 (other Esx-related and pyrimidine synthesis proteins are not shown). Data represent the mean ± s.d. for proteins in the indicated category, n = 3 technical replicates. *p≤0.05, one sample t test comparing mean with value 0. B) EsrG1 T263 phosphorylation in *S. meyeri* W712 wild-type grown in mono-cultures (Control) or co-cultures with TM7-008 or TM7- 074, based on MaxQuant phosphor (STY) analysis of whole cell proteome datasets. The line indicates the mean. n = 3 technical replicates. Values are raw MS1 intensities (arbitrary units). C) Unrooted phylogenetic tree depicting the distribution of Esr pathways across the class Actinomycetes. Each leaf represents a taxonomic family, colored according to the number of Esr pathways identified among members of that family. Orders containing greater than one family are highlighted in grey. An additional blue circle denotes families containing species that have been experimentally shown to support the growth of Saccharibacteria. Tree is based on an concatenated alignment of 120 conserved proteins, derived from the Genome Taxonomy Database^77^. D) Schematic depicting representative examples of genetically linked Esr and Esx gene clusters. Genes are colored according to functional modules as in Figure 2A. E) Schematic depicting the domain architectures of diverse EsrG proteins and their relative frequencies. The experimentally determined phosphorylation sites on EsrG proteins from *S. odontolytica* and *S. meyeri* are depicted as yellow circles. F) Relative levels (SC ratio) of the indicated proteins in cell-free supernatant of the indicated So F0309 strains, normalized to wild-type values. Data represent the mean ± s.d, n = 3 technical replicates. G) Fold-change in levels of TM7-008 after 40 hours of growth on the indicated So F0309 strains. Data represent mean ± SD (n = 3 biological replicates). * p≤0.05, one-way ANOVA with Šidák correction for multiple comparisons to the mean of the wild-type strain; H) Model depicting the potential modes of Esr pathway function, with Esr components colored to reflect relative sequence diversity. The sequence diversity of the linker domain of EsrA, which can be comprised of different classes of repeats, could not be readily compared across proteins and is thus colored grey. In both Esr1-like (left) and Esr2-like pathways (right), sensing of diverse threat signals via the sequence-divergent β-propeller domains of EsrA proteins activates a signaling cascade that proceeds through conserved MoxR and vWA proteins and a kinase/phosphatase pair to affect the phosphorylation state of a divergent FHA protein. Upon phosphorylation, the variable domain architectures of these FHA proteins permit a variety of responses, including transcription of defensive genes through direct (zinc finger domain-mediated) and indirect mechanisms (left) and post- translational activation of defensive pathways such as the Esx secretion system (right). Alongside canonical EsxA-EsxB dimers (grey), these Esx systems secrete AesB-like effectors (green) that promote defense against the threat imposed by Saccharibacteria. PPL1, pseudophosphatase-like; Zn, zinc finger.

Motivated by the apparent generality of the Esr pathway and the response it controls across multiple species, we performed a search for the pathway in publicly available bacterial genomes. We found that phylogenetically, the Esr pathway is effectively confined to bacteria in the class Actinomycetes, with particular prevalence across the orders Actinomycetales and Propionibacteriales (Figure 4C, Supplemental Table 3). Intriguingly, these orders include all species experimentally demonstrated to support growth of one or more Saccharibacterial strains, supporting a broader link between Esr and defense against Patescibacteria^17,24,50–52^. Outside of Actinomycetales and Propionibacteriales, Esr distribution is patchy and confined to a limited number of species within the Mycobacteriales and Streptomycetales, suggesting it may have been horizontally acquired in these groups.

Our comprehensive search identified a strong genomic link between the Esr and Esx pathways; 40% of the 3502 Esr pathways we identified are encoded immediately adjacent to *esx* genes (Figure 4D, Supplemental Table 3). This is a conservative estimate, as our criteria for Esr identification required a complete complement of So F0309 Esr1 orthologs. While this genetic linkage provides evidence for the functional association of the pathways, we additionally identified numerous instances in which the critical signal output protein, EsrG, is translationally fused to the essential Esx ATPase EccC (Figure 4E). Together, these observations prompted us to investigate whether the Esx-linked Esr2 pathway of So F0309 regulates Esx activity in a manner distinct from the transcriptional control imposed by Esr1. Semi-quantitative mass spectrometry analysis of culture supernatant deriving from So F0309 Δ*esrA2* revealed that the levels of the EsxA and EsxB approached those found in wild-type culture supernatant (Figure 4F). However, AesA and AesB were undetectable in the So F0309 Δ*esrA2* secretome despite intracellular levels approximating that of the wild-type control (Supplemental Table 2). This finding, together with the observation that Esx protein levels are not significantly altered in the Δ*esrA2* background (Supplemental Figure 8A, Supplemental Table 2), suggests that the Esr2 pathway acts posttranslationally to influence the secretion of a subset of Esx substrates. A unique property of AesA and AesB is that they appear to contain a variant of bipartite signal known to mediate secretion of other Esx substrates (FXG between α-helices 1 and 2 in AesB and a proximal aspartate near the N-terminus of AesA in place of the canonical WXG or LXG and YxxxD/E motifs), which could provide a biochemical basis for substrate selectivity in Esr-regulated systems (Supplemental Figure 8F)^25^. In support of the specialized role of Esr2 in regulating Esx activity, we find that TM7-008 achieves significantly higher growth yields on So F0309 Δ*esrA2* compared to the wild-type strain (Figure 4G); So F0309 Δ*esr1* was not tested in this assay due to the confounding effects of modifying its large regulon. Together, these data provide a genetic and functional link between a dedicated threat sensing pathway found across Actinobacteria and a secretion system capable of inhibiting epibiont growth.

While our results clearly point to a link between the Esr and Esx pathways and a response to epbiotic bacteria, we have yet to establish how Esr systems sense epibionts, nor have we determined how EsrG phosphorylation leads to changes in transcription. If Esr is beneficial to host bacteria under ecologically relevant conditions, then we would anticipate an arms race dynamic promoting diversification of the protein responsible for threat detection and its corresponding ligand on the epibiont. Indeed, we find that the surface-localized β-propeller domain of EsrA is among the most sequence diverse elements across Esr pathways and we have identified a Saccharibacteria strain that evades detection by Esr (Figure 3G, Figure 4H). Studies of other threat sensing modules often find that the output of the pathway is also diversified^5,47^. Our global analysis of Esr pathways found that while EsrG proteins consistently contain an FHA domain, this is linked to at least 13 distinct domain configurations (Figure 4E, Supplemental Table 3). Prominent among these are variants of the nucleic acid-interacting zinc finger domain, suggesting that EsrG could be directly involved in modulating transcription in these cases. Our bioinformatic analyses thus point to the Esr pathway serving as an adaptable, functionally plastic mechanism with the potential to promote Actinobacterial survival in the face of a wide range of bacterial antagonists.

## Discussion

In this study, we discover a threat-sensing pathway, termed Esr, that mediates an adaptive response to epibiont colonization and demonstrate that one target of this response, the Esx secretion system, inhibits epibiont proliferation. The epibiont-inducible nature of the system argues Esr is an intrinsically defensive pathway. This is in-line with other intermicrobial antagonism response systems, including the Gac/Rsm-regulated unified antagonism response and defense (GUARD) system found in Pseudomonads^5–7^. For several characterized factors within the GUARD regulon, it is clear why linking their production (or removal) to the presence of an immediate threat is beneficial. For example, GUARD induces the metabolically costly antibacterial T6SS and potentially lethal abortive phage defense systems^4,53^. Mirroring GUARD, Esr induces an Esx secretion system, which we suggest serves a specialized role in antibacterial antagonism akin to the T6SS. The cell surface modifications induced by Esr may also be detrimental in the absence of an epibiontic threat. Zhong *et al.* found that the predicted cell wall-associated polysaccharide biosynthesis genes induced by Esr confer susceptibility to lytic phage LC001 in the related host strain, So XH001^23^. One striking difference between Esr and GUARD, however, is that GUARD responds to pseudomonad lysis and is thereby activated in response to a wide variety of threats, while the Esr pathway can discriminate between closely related Patescibacteria. This observation suggests that Esr is not broader danger sensing pathway like GUARD, but rather is specialized to sense and defend against particular epibionts.

Our discovery and initial characterization of the Esr pathway provides a new window into the ecological relationship between Patescibacteria and their hosts. In total, we find our data support one of antagonism between these bacteria. The strong genomic and functional links between the pathway and Esx, taken together with our experimental findings demonstrating that Esx limits Patescibacterial proliferation, suggests that unchecked growth of the epibiont poses a burden to its host. This appears not to be limited to the So F0309–TM7-008 interaction we focused our experimental work on, as Esr is encoded across Actinomycetales and it remains highly associated with Esx in these additional organisms. Furthermore, extensive bioinformatic analyses conducted by Aravind and colleagues indicate that Esr components have evolved to serve at the interface of biological conflicts^47^. A number of closely related pathways identified by these authors, which consist of sensing modules linked to the MoxR–vWA protein pair, additionally bear effector domains implicated directly in defense against phage and other foreign DNA elements.

We have identified a previously undescribed class of Esx substrates found in Actinomycetales and Propionibacteriales, and show that one of these proteins, AesB of So- F0309, limits proliferation of the epibiont TM7-008. We additionally show that secretion of the AesA–AesB complex requires the Esr2 pathway encoded adjacent to the Esx secretion system, whereas this pathway is dispensable for EsxA and EsxB export. Substrate heirarchy is common in Esx secretion systems, though outside of the Actinomycetales it is believed to stem from early substrates triggering rearrangement of the apparatus through the ATPase domains of EccC^33,34^. The EsrG2 protein that likely mediates the output of Esr2 signaling contains a C-terminal FHA domain. In Firmicutes, tandem FHA domains located at the N-terminus of EccC are required for Esx apparatus assembly^54,55^. Together, these observations suggest a model in which the switch between early and late effector translocation in So F0309 and related Actinobacteria relies on modulation of EccC activity by the FHA domain of EsrG2.

Relative to their prevalence across ecosystems, Patescibacteria–host interactions remain significantly understudied. Slow development of the field initially stemmed from limitations in sequence-based methods for surveying bacterial diversity, whereas current challenges include identifying physiological hosts to support Patescibacteria cultivation, and limited tools inherent to working with non-model organisms; the latter amplified in a system requiring co-cultivation of two such organisms^16^. While our current study highlights the need to interrogate Patescibacteria–host interactions in order to understand the full complement of evolutionary pressures driving bacterial molecular innovations, the questions that remain underscore the challenges associated with a system with few existing tools and little underlying literature. Areas requiring further investigation include identifying the epibiont ligand that interacts with EsrA1, defining the mechanism by which EsrG1 phosphorylation triggers changes in transcription, understanding how the Esr2 pathway is activated and how this influences Esx secretion, and elucidating the mechanism by which AesB restricts epibiont proliferation. Furthermore, in light of our finding that hosts actively antagonize epibionts that colonize them, it stands to reason that Patescibacteria have evolved mechanisms to subvert Esr signaling or evade Esx targeting. Considering the ubiquity of this interaction and the extent of uncharacterized genes in these non- model organisms, the study of these evolutionary conflicts promises to reveal fundamentally new facets of bacterial biology.

## Methods

### Bacterial strains and culture conditions

A complete list of bacterial strains used in this study can be found in Supplementary Table 4. All strains generated in this study are available upon request from the corresponding author. *Escherichia coli* strain DH5α was used for plasmid cloning and was grown in Lysogeny broth (LB) media at 37°C with shaking or on LB agar plates. Unless otherwise stated, *Schaalia odontolytica* F0309 and *S. meyeri* W712 mono-cultures and co-cultures with *Nanosynbacter* sp. TM7-008 or *Nanosynbacter* sp. TM7-074 were grown statically at 37°C under microaerophilic condition using GasPak EZ Campy Container System Sachets (BD 260680) in Brain Heart Infusion (BHI) media or on BHI agar plates. *Southlakia epibionticum* ML1 (Se)–*Actinomyces israelii* F0345 (Ai) co-cultures were grown in TSY media (30 g/L tryptic soy and 5 g/L yeast extract) at 37°C in ambient air enriched with 5% CO_2_ with shaking. Media were supplemented with antibiotics when needed at the following concentrations: kanamycin (kan; 50 µg ml^-1^ for *E. coli*, 100 µg ml^-1^ for *S. odontolytica* and *S. meyeri*), hygromycin (hyg; 150 µg/mL). *S. odontolytica* and *S. meyeri* mono-cultures and co-cultures with Saccharibacteria were stored at −80°C in BHI supplemented with 18% (v/v) glycerol.

### Plasmid construction

Plasmids, synthetic DNA fragments and primers used in this work are provided in Supplementary Table 4. Plasmids generated in this study are available upon request from the corresponding author. Synthetic DNA fragments were obtained from Twist Bioscience and primers from Integrated DNA Technologies. All plasmid constructs were designed using Geneious Prime, generated using Gibson assembly^56^, and confirmed by sequencing.

### *S. odontolytica* F0309 genome sequencing

The publicly available genome sequence for *S. odontolytica* F0309 (GCA_000163415.1) consists of multiple unassembled contigs, complicating transcriptomic and proteomic analyses. To obtain a closed reference genome to be employed in subsequent analyses, we sequenced the genome of *S. odontolytica* F0309 wild-type strain using the Standard Bacterial Genome Sequencing with Extraction service by Plasmidsaurus (Eugene, OR, USA). Briefly, cells were submitted in 1x DNA/RNA Shield (Zymo Research), and amplification-free libraries were sequenced on an Oxford Nanopore flow cell. Reads were filtered with Filtlong v0.2.1, assembled with Flye v2.9.1 and polished with Medaka v1.8.0. Annotation was performed with Bakta v1.11. This sequence and annotation will be deposited in Genbank.

### Construction of genetically modified *S. odontolytica* strains

*S. odontolytica* mutants were generated via allelic exchange using the newly generated plasmid pLG202, which has a modified alpha subunit of phenylalanyl-tRNA synthetase gene that enables counter selection with 4-chloro-DL-phenylalanine, and a kanamycin resistance cassette derived from pCWU3^57^ for selection in *Schaalia* species. Briefly, DNA fragments that include homology arms of 900–1000 bp to facilitate chromosomal integration by homologous recombination were cloned into pLG202 using Gibson assembly^56^. For gene complementation in So F0309, DNA fragments were inserted into a neutral site (NS; located at the intergenic region between OBNOAD_02079 and OBNOAD_02080 in *S. odontolytica* F0309 genome) using pLG202. Electrocompetent *S. odontolytica* cells were prepared as previously described, with modifications noted below^58,59^. Log-phase *S. odontolytica* cultures were mixed with prewarmed BHI + 20% glycine to for a final concentrations of 7.5% glycine and incubated under microaerophilic condition with shaking at 37°C for 1 h. Cultures were then incubated in ice slurry for 30 min with periodical swirling and cells were kept at 4°C from this point on. Cells were pelleted at 3,000 rcf for 15 min at 4°C, washed with ice-cold 10% glycerol for 4 times, and resuspended in a small volume of 10% glycerol. After aliquoting, cells were flash frozen in dry ice/ethanol and store at −80°C. Plasmids were electroporated into *S. odontolytica* electrocompetent cells using the following parameters: 2.25 kV, 400 Ω, and 25 µF with 1 mm cuvettes. After pulsing, cells were mixed immediately with prewarmed BHI media and recovered for 4 h under microaerophilic condition statically at 37°C. Cells were then concentrated by centrifugation at 4,000 rcf for 5 min and plated on BHI plates supplemented with 100 µg/mL kan. The resulting colonies were cultured in BHI media (no kan) to saturation. The cultures were then diluted 1:1000 in BHI + 10 mM DL-4-chloro-phenylalanine and grown for 2–3 days and streaked on BHI plates (no kan) for single colonies. Isolated colonies were screened for kan sensitivity by patching on BHI + kan100 and BHI plates, and kan-sensitive colonies were genotyped by PCR.

### Construction of genetically modified Saccharibacteria

TM7-008–sfGFP mutant was constructed essentially as described previously with the modifications described below^60^. Overnight *S. odontolytica* F0309 cultures were diluted to OD = 0.4 with BHI and mixed with purified TM7-008 and DNA fragments for inserting *sfgfp* (under the control of *tuf* promoter from *Nanosynbacter lyticus* TM7x) and *hph* (hygromycin resistance gene; under the control of *rpsJ* promoter from *N. lyticus* TM7x) into a neutral site (NS; located at the intergenic region between LRM48_003185 and LRM48_003190 based on NCBI accession CP157587.1). The mixtures were incubated under microaerophilic condition statically at 37°C for 6 h and diluted ∼3 fold in BHI with hyg. Cultures (P0) were incubated under microaerophilic condition statically at 37°C for ∼24 h and passaged 1:10 into BHI supplemented with hyg (P1). Cultures were subsequently passaged every ∼24 h in the same way to P4. After ∼24 h of growth, P4 cultures were determined to have high number of GFP-positive TM7-008 by flow cytometry and were streaked on BHI plates. The resulting single colonies were inoculated into BHI media and after growth, screened by flow cytometry for co-cultures containing GFP-positive TM7-008. The TM7-008–sfGFP–*S. odontolytica* F0309 co-cultures were passaged several times and filtered using 5 µm and 0.2 µm filters sequentially, and the filtrates were used to infect *S. meyeri*- W712, a strain that supports particularly robust growth of TM7-008. After one day of growth, the TM7-008–sfGFP*–S. meyeri-*W712 co-cultures were used to generate glycerol stocks and stored at −80°C.

### Purification of Saccharibacteria

To create purified suspensions of TM7-008 and TM7-074, co-cultures of TM7-008–*S. meyeri* W712 or TM7-074–*S. meyeri* W712 grown in BHI were amended with L-arginine (10 mM final concentration) and centrifuged at 3,500 rcf for 10 min at room temperature to pellet host cells. Supernatant containing Saccharibacteria was then sequentially filtered with 5 µm, 1.25 µm, and 0.45 µm filters. Filtrate was centrifuged at 80,000 rcf for 25 min at room temperature to pellet Saccharibacteria. The resulting pellet was resuspended in a small volume of BHI with 18% (v/v) glycerol and 10 mM L-arginine, flash frozen in a dry ice and ethanol mixture, and stored at −80°C. To create purified suspensions for *Se*, co-cultures of *Se* and *Ai* F0345 grown in TSY media were filtered with 0.45 µm filters, and filtrate was spun down at 15,000 rcf at 4°C for 4 hours. Pellets were resuspended in TSY supplemented with 10% (v/v) dimethylsulfoxide (DMSO), flash frozen in a dry ice and ethanol mixture, and stored at −80°C.

### Saccharibacteria host compatibility testing

Purified populations of *S. epibionticum*, *Nanosynbacter* sp. TM7-008, and *Nanosynbacter* sp.TM7-074 were prepared as described above. To assess growth of each Saccharibacteria strain on *S. odontolyltica* F0309, triplicate stationary phase cultures of the host strain were diluted to OD_600_ = 0.05 in 4 mL BHI and incubated for 4 hours at 37°C. Cultures were subsequently diluted to OD_600_ = 0.01 in 4 mL BHI, infected with each purified Saccharibacteria strain at an MOI of 10, and incubated at 37°C under microaerophilic conditions for an additional two hours. Saccharibacteria*/S. odontolytica* F0309 co-cultures were subsequently diluted 3200-fold in BHI and returned to 37°C microaerophilic conditions for 48 hours, then 1 mL aliquots were collected for bacterial quantification by qPCR analysis. To assess growth of Saccharibacteria on the host strains *S. meyeri* W712 and *A. israelii* F0345, triplicate stationary phase cultures of each host were diluted in 1 mL BHI to OD_600_ = 0.2 and infected with each purified Saccharibacteria strain at an MOI of 0.2. 100 µL of each Saccharibacteria/host co-culture was collected by centrifugation at 21 000xg for 30 minutes and stored at −20°C for qPCR analysis of the starting populations. Co-cultures were incubated at 37°C under microaerophilic conditions for 48 hours (*A. israelii*) or 72 hours (*S. meyeri*), and 100 µL aliquots were collected for qPCR analysis.

### Transcriptomics

Overnight *S. odontolytica* F0309 cultures were back-diluted to OD = 0.3 and incubated under microaerophilic condition statically at 37°C for 4 h. Cultures were then diluted and mixed with either purified Saccharibacteria at MOI∼35 or the same diluent used in generating Saccharibacteria stocks, with a final *S. odontolytica* F0309 OD of 0.5. These mixtures were incubated under microaerophilic condition statically at 37°C for 2 h and then mixed with 2 volumes of RNAprotect Bacteria Reagent (Qiagen). After centrifugation at 6,000 rcf for 15 min at room temperature, supernatants were removed and pellets were flash frozen in liquid nitrogen and stored at −80°C. RNA was either extracted in-house following RNAprotect protocol for enzymatic lysis, proteinase K digestion and mechanical disruption of bacteria followed by purification of total RNA from bacterial lysate using the RNeasy Mini Kit (Qiagen), or extracted by SeqCenter (Pittsburgh, Pennsylvania) with ZymoBIOMICS^TM^ Quick-RNA Miniprep Kit (Zymo Research, R1055). After RNA extraction, samples were processed and sequenced by SeqCenter. Per SeqCenter protocol, samples were treated with DNase (RNase free; Invitrogen), and library preparation was performed using Stranded Total RNA Prep Ligation with Ribo-Zero Plus kit for ribosomal RNA depletion (Illumina) and 10 bp unique dual indices (UDI). Sequencing was carried out on a NovaSeq X Plus, producing paired end 150 bp reads. Demultiplexing, quality control, and adapter trimming was performed with bcl-convert (Illumina). Read mapping was performed with HISAT21^61^, and read quantification was performed using featureCounts in Subread^62^. Read counts loaded into R (https://www.r-project.org/) were normalized using the Trimmed Mean of M values algorithm in edgeR, and differential expression analysis was performed using glmQLFTest in edgeR^63^. Transcriptomic RNA-seq data are in the process of being depositing in the NCBI Sequence Read Archive under Bioproject SUB16465414.

### Whole cell proteomics

For measuring proteomic changes in host bacteria (*S. odontolytica* or *S. meyeri*) with or without Saccharibacteria, log phase host bacteria were diluted and mixed with either purified Saccharibacteria at MOI ≥ 15 or the same diluent used in generating Saccharibacteria stocks, with a final host bacterium OD of 0.1. The resulting cultures were incubated under microaerophilic condition with shaking at 37°C for 7 h. After incubation, cultures were centrifuged at 5,000 rcf for 10 min at room temperature. Pellets were resuspended in PBS and cells were centrifuged again. The resulting pellets were flash frozen in liquid nitrogen and stored at −80°C. To prepare samples for mass spectrometry, 250 µL lysis buffer (8 M urea, 75 mM NaCl, 50 mM Tris-HCl, pH 8.2) was added to previously-frozen cells and cells were lysed by sonication in 1.5 mL microcentrifuge tubes. After spinning at 20,000 rcf for 10 min, the protein concentration of the resulting supernatant measured using the Pierce™ BCA Protein Assay Kit (Thermo Fisher Scientific). Meanwhile, 200 µL of the protein solutions were transferred to new 1.5 mL microcentrifuge tubes and incubated with 5 mM dithiothreitol (DTT) at 56°C for 25 min for protein reduction. After cooling to room temperature, iodoacetamide was added to 14 mM and the samples were incubated in the dark at room temperature for 30 min for alkylation. The reaction was quenched by adding additional 5 mM DTT and incubating 15 min in the dark at room temperature. Based on BCA assay, the same amount of protein from each sample (within a single experiment) was taken to a new tube and was diluted 5-fold with 25 mM Tris-HCl (pH 8.2) to reduce the urea concentration to 1.6 M and 1 mM calcium chloride was then added. Samples were mixed with trypsin (Promega) and incubated at 37°C for 17 h. Digestion was halted by adding 0.4% trifluoroacetic acid (TFA). Peptides were purified using either C18 columns (BioPureSPN MACRO PROTO^TM^300 C18, 50-200 µL loading, 35-350 µg capacity, The Nest Group HMM S18V) or in-house-prepared stop-and-go-extraction tips (StageTips)^64^ embedded with Empore™ styrene divinyl benzene (SDB-RPS) extraction disks (Sigma), or both C18 columns and StageTips sequentially. For peptide purification with C18 columns: C18 columns were charged by washing with 250 µL of 100% acetonitrile (ACN), followed by two washes with 250 µL water before loading samples. Bound samples were washed twice with 250 µL of 5% ACN + 0.1% TFA and eluted with 250 µL of 80% ACN + 25 mM formic acid. Eluted peptide samples were then dried in a SpeedVac^TM^ vaccum concentrator (Thermo Fisher Scientific). For StageTips purification: StageTips were conditioned sequentially with 100 µL of 100% methanol, 50 µL of 100% ACN, 50 µL of 75% ACN + 5% ammonium hydroxide, 50 µL of 75% ACN + 0.5% acetic acid, and 100 µL of 0.1% TFA. Peptides were loaded onto conditioned StageTips and washed sequentially with 100 µL of 0.1% TFA, 100 µL of 75% ACN + 0.5% acetic acid, 50 µL of 0.5% acetic acid. Peptides were then eluted with 75 µL of 75% ACN + 5% ammonium hydroxide, and eluted peptide samples were dried in a SpeedVac^TM^ vaccum concentrator. Dried peptides were resuspended in 5% ACN and 5% formic acid and analyzed by LC-MS/MS on am Orbitrap Fusion Lumos mass spectrometer (Thermo Scientific) as previously described^7^. Mass spectrometry data were analyzed using MaxQuant^65^. For plotting protein differential expression graphs based on whole cell proteome datasets, missing values in normalized LFQ intensity were imputed from a normal distribution of log-transformed data defined by a mean at (data median – 1.8 x std. dev) and a standard deviation of (data std. dev ∗ 0.5), akin to previously described methods^66^.

### Secretomics

All cultures in secretomic experiments were grown in tryptic soy broth (TSB) media to reduce background signal in LC-MS/MS caused by BHI media. Overnight *S. odontolytica* F0309 cultures were back-diluted to OD = 0.1 and incubated under microaerophilic condition statically at 37°C for 7 h. Cultures were then centrifugated at 6,000 rcf for 20 min at room temperature and the supernatant fraction was removed. The pellet fraction was washed once with PBS, centrifuged again, and the resulting pellets were flash frozen in liquid nitrogen, stored at −80°C, and later processed for and analyzed with mass spectrometry as described above for whole cell proteomics. The supernatant fraction was filtered with a 0.2 µm filter to remove any remaining cells. Protease inhibitors (Roche 11873580001 cOmplete, EDTA-free Protease Inhibitor Cocktail) were added and supernatant samples were concentrated to a smaller volume using 3 kDa MWCO Amicon Ultra Centrifugal Filter (Millipore). The concentrated supernatant samples (1.75 mL each) were dialyzed in 3.5 K MWCO Slide-A-Lyzers (Thermo Scientific) against PBS at 4°C with stirring (3 rounds of dialysis each with 1 L PBS). After dialysis, supernatant samples were mixed with trichloroacetic acid (TCA) to a final concentration of 20% and incubated on ice for 30 min to precipitate proteins. After centrifugation at 13,500 rcf for 10 min at 4°C, the pellet was washed once with 20% TCA and twice with 100% acetone. The dried protein samples were resuspended in lysis buffer (8 M urea, 75 mM NaCl, 50 mM Tris-HCl, pH 8.2) followed by reduction, alkylation, trypsin digestion, peptide purification with StageTips, and mass spectrometry analysis as described above for whole cell proteomics.

### Co-immoprecipitation assays

*S. odontolytica* cultures encoding VSV-G-tagged alleles of *aesA* and *aesB* at the native chromosomal loci were subcultured to an OD_600_ of 0.005 in 100 mL BHI and incubated shaking at 150 rpm at 37°C under microaerophilic conditions for 8 hours. Cultures were divided into two technical replicates and collected by centrifugation at 8000xg for 20 mins. Pellets were washed with 1 mL wash buffer (150 mM NaCl, 20 mM Tris HCl pH 7.5, 2% (v/v) glycerol, and 2 mM beta-mercaptoethanol) and collected by centrifugation at 8000xg for 5 minutes. Pellets were stored at −80°C overnight.

To immunoprecipitate VSV-G-tagged proteins and binding partners, cells were resuspended in 0.5 mL lysis buffer (150 mM NaCl, 20 mM Tris HCl pH 7.5, 2% (v/v) glycerol, and 2 mM beta-mercaptoethanol, 0.1% Triton-X 100, 0.1% IGEPAL, 25 U/mL benzonase) and lysed on ice by 6 rounds of 15 second sonication pulses at 35% power using a Q125 ultrasonic processor equipped with a 1/8” microtip (QSonica). Lysates were clarified by centrifugation at 21 000xg at 4°C. 5 µg rabbit polyclonal anti-VSV-G antibody (Sigma) was added to each sample and samples were incubated at 20°C in a thermal mixer shaking at 500 rpm for 1.5 hours. Samples were subsequently added to 25 µL protein A/G magnetic beads (ThermoFisher) and incubated in a thermal mixer shaking at 500 rpm at 20°C for an additional 2 hours. Beads were magnetically separated from the bulk lysate, washed 5x with 1 mL wash buffer, washed twice with 1 mL 20 mM ammonium bicarbonate, and resuspended in 30 µL 20 mM ammonium bicarbonate. Protein A/G agarose beads and bound proteins were treated with 10 µL 10 ng/µL sequencing-grade trypsin (Promega) for 16 hours at 37°C shaking at 300 rpm. Beads were washed twice with 50 µL 20 mM ammonium bicarbonate and each wash was collected and combined as the peptide fraction. Peptides were reduced with 1 mM tris(2-carboxyethyl) phosphine hydrochloride at 37 °C for 1 h, followed by alkylation with 10 mM iodoacetamide at room temperature in the dark for 30 min. The alkylation reaction was quenched with 6 mM dithiothreitol at room temperature in the dark for 15 min. Samples were desalted using the same method as described above for whole cell proteome samples.

### Measurement of Saccharibacteria proliferation

To compare proliferation of Saccharibacteria (TM7-008 or TM7074) during co-culture with different *S. odontolytica* strains, stationary phase cultures of *S. odontolytica* strains to be tested were subcultured in triplicate to an OD_600_ of 0.05 in 4 mL BHI and incubated statically at 37°C under microaerophilic conditions for 4 hours. Cultures were subsequently diluted to OD_600_ of 0.01 in 2 mL BHI and 8 x 10^7^ purified Saccharibacteria were added to achieve a multiplicity of infection of 10. Cultures were returned to 37°C and microaerophilic conditions for an additional 2 hours to allow for Saccharibacteria attachment before being diluted 3200-fold into 4 mL fresh BHI. A 0.5 mL aliquot of undiluted Saccharibacteria/host mixture (representing the initial timepoint) was collected by centrifugation at 21 000xg for 30 minutes and stored at −20°C for qPCR analysis. Diluted Saccharibacteria/host mixtures were incubated for 40 hours at 37°C under microaerophilic conditions, after which 1 mL of mixture was collected by centrifugation at 21 000xg for 30 minutes and stored at −20°C for qPCR analysis. For experiments involving *S. odontolytica* strains encoding genes of interest under the control of theophylline-responsive riboswitches (i.e., *aesA*, *aesB* complemented *in trans*), theophylline was included in all cultures at 0.2 mM.

### qPCR analysis of Saccharibacteria and host proliferation

qPCR was employed to quantify the proliferation of Saccharibacteria and host populations in co-cultures. Genomic DNA was isolated from frozen cell pellets using 100 µL InstaGene matrix (BioRad catalogue 7326030) according to the manufacturers instructions. Quantitative PCR (qPCR) was performed in 20 µL reactions containing 300 nM each primer, 4 µL template, and 1x SsoAdvanced Universal SYBR Green Supermix (BioRad catalogue 1725272). Primer sequences are provided in Supplemental Table 4. Thermocycling conditions were 95°C for 5 minutes, followed by 35 cycles of 95°C for 20 s, 60°C for 30 s, and a fluorescent reading. Target sequence abundance was determined by comparison to standard curve reactions performed in parallel with each assay. Standard curve templates were prepared from genomic DNA isolated from host cultures or purified Saccharibacteria using the Wizard HMW DNA extraction kit (Promega catalogue A2920) and quantified by Qubit (ThermoFisher).

### Measurement of host proliferation during Saccharibacteria infection

A colony-forming unit (CFU)-based assay was used to determine the effect of Saccharibacteria infection on *S. odontolytica* proliferation. Stationary phase cultures of *S. odontolytica* strains were subcultured in triplicate in 4 mL BHI to an OD_600_ of 0.05 and incubated at 37°C under microaerophilic conditions for 4 hours. Cultures were subsequently diluted to OD_600_ of 0.01 in 1 mL BHI and 4 x 10^8^ purified Saccharibacteria were added to achieve a multiplicity of infection of 100. A set of replicate cultures was prepared without Saccharibacteria to serve as matched uninfected controls. Saccharibacteria/host co-cultures were incubated statically for 16 hours at 37°C under microaerophilic conditions. To quantify host CFUs at the initial timepoint, 100 µL of Saccharibacteria/host co-culture was collected and diluted in 10-fold series in a 96 well microtiter plate, and 7 µL of the 10^0^ to 10^-5^ dilutions were spotted onto BHI containing 1.5% agar. To quantify host CFUs at the 16-hour timepoint, 100 µL of Saccharibacteria/host co-culture was collected and diluted in 10-fold series in a 96 well microtiter plate, and 4 µL of the 10^-1^ to 10^-6^ dilutions were spotted onto BHI containing 1.5% agar. Plates were incubated at 37°C under microaerophilic conditions for 36 hours before colonies were counted. To determine infection-dependent growth inhibition, the growth of each culture was calculated as the ratio of final/initial CFU/mL, and the growth of each uninfected control culture was divided by the growth of the matched Saccharibacteria-infected culture.

### Flow cytometric determination of Saccharibacteria infection burden

A flow cytometry-based assay was used to quantify the burden of Saccharibacteria infection with or without significant dilution. Stationary phase cultures of *S. odontolytica* were subcultured in triplicate in 4 mL BHI to an OD_600_ of 0.05 and incubated at 37°C under microaerophilic conditions for 4 hours. Cultures were subsequently diluted to an OD_600_ of 0.01 in 2 mL BHI, and 8 x 10^7^ purified TM7-008 constitutively expressing sfGFP from a neutral chromosomal site was added to achieve a multiplicity of infection of 10. An additional set of matched uninfected control cultures were also prepared to assess background GFP levels. After 2 hours of incubation at 37°C under microaerophilic conditions, 12 µL of co-culture or matched uninfected control culture was collected and diluted 3200-fold into a final volume of 4 mL BHI to serve as the low infection condition. Both the diluted and non-diluted cultures (representing the high infection condition) and matched uninfected controls were returned to 37°C under microaerophilic conditions for 16 hours. Cultures were diluted 20-fold in filtered PBS (pH 7.4) and analyzed on a BD Symphony A1 flow cytometer equipped with a small particle detector. Host cells were first gated by forward and side scatter, and infected cells were identified by GFP fluorescence above the background level of the uninfected controls.

### Bioinformatics analysis

#### Identification of candidate Esx substrates of S. odontolytica

Candidate Esx substrates were identified in *S. odontolytica* on the basis of their predicted structural similarity to characterized substrates. We initially searched for candidate substrates by using AlphaFold3 to predict structures for each protein encoded within the Esx secretion system (OBNOAD_00322-00338)^67^. Those proteins sharing structural similarity with Esx substrates were identified by Foldseek ^68^. To identify distally encoded candidate substrates, we used BLASTp to search for homologs of the candidates we identified withing the Esx gene cluster (OBNOAD_00323: AesA, OBNOAD_00324:AesB, OBNOAD_00330: EsxA, OBNOAD_00331: EsxB, OBNOAD_00337 and OBNOAD_003338), which led to the identification of AesB homologs OBNOAD_01965, OBNOAD_00733, and OBNOAD_00094. Each of these proteins is encoded adjacent to a predicted EsxA/B-like protein, and BLASTp searches of these led to identification of an additional candidate substrate pair, OBNOAD_01274-1275. The remainder of the candidate substrates were identified by modeling the structures of Esr-regulated proteins with AlphaFold3 and comparing these to characterized Esx substrates through Foldseek searches.

#### Phylogenetics and conserved sequence analysis of AesB

Homologs of AesB were identified by performing iterarted PSI-BLAST searches of the Clustered NR database until no new sequences sharing homology across at least 80% of the protein were identified (four rounds)^69^. This led to the identification of 181 homologous sequence clusters. Proteins representative of each cluster were aligned using MUSCLE in Geneious Prime, and positions containing >15% gaps were stripped^70^. A phylogeny based on this alignment was constructed using the RAxML implementation in Geneious Prime^71^. Through the phylogenetic analysis, a clade containing 103 AesB homologs with a conserved charged region in their C-terminus was identified, and these proteins were re-aligned using MUSCLE to identified conserved sequence motifs. Intrinsically disordered regions in AesB were identified using IUPred3^72^.

#### Identification of Esr pathways and their genetic context

To comprehensively identify genomic loci containing esr pathways, we searched for homologs of components from the representative operons from *S. odontolytica* F0309 using iterative PSI-BLAST^69^ searches against the NCBI non-redundant (nr) protein database until convergence, with an E-value cutoff of 0.005. For each identified homolog, we retrieved its genomic neighborhood, including 10 genes upstream and 10 genes downstream, from the NCBI GenBank database^73^. All neighboring proteins were clustered based on sequence similarity using BLASTCLUST, a BLAST score-based single-linkage clustering method (https://ftp.ncbi.nih.gov/blast/documents/blastclust.html). Protein clusters were subsequently annotated based on their domain architectures using HMMSCAN^74^ against the Pfam database^75^ and our in-house custom HMM profile database. Signal peptides and transmembrane regions were predicted using Phobius^76^.

## Acknowledgements

We thank Judit Villen, Ricard Rodriguez-Mias and Jimmy Eng for help with mass spectrometry data acquisition and analysis, and Jeff McLean, Xuesong He and members of the Bor and Mougous laboratories for helpful discussions. Support for this work was provided by a Project Grant from the Canadian Institutes of Health Research (PJT-206152 to J.C.W.), the National Institutes of Health (DE031274 to B.B.), the Saint Louis University President’s Research Fund (D.Z.) and the University of Washington’s Proteomics Resource (UWPR95794). J.C. is supported by a Canada Graduate Scholarship – Doctoral and a Michael Smith Foreign Study Supplement from the Natural Sciences and Engineering Research Council of Canada. J.D.M. is an HHMI Investigator and holds the John F. Enders Professorship in Microbial Pathogenesis at Yale University.

## Supplemental Figure Legends

**Supplemental Figure 1.**
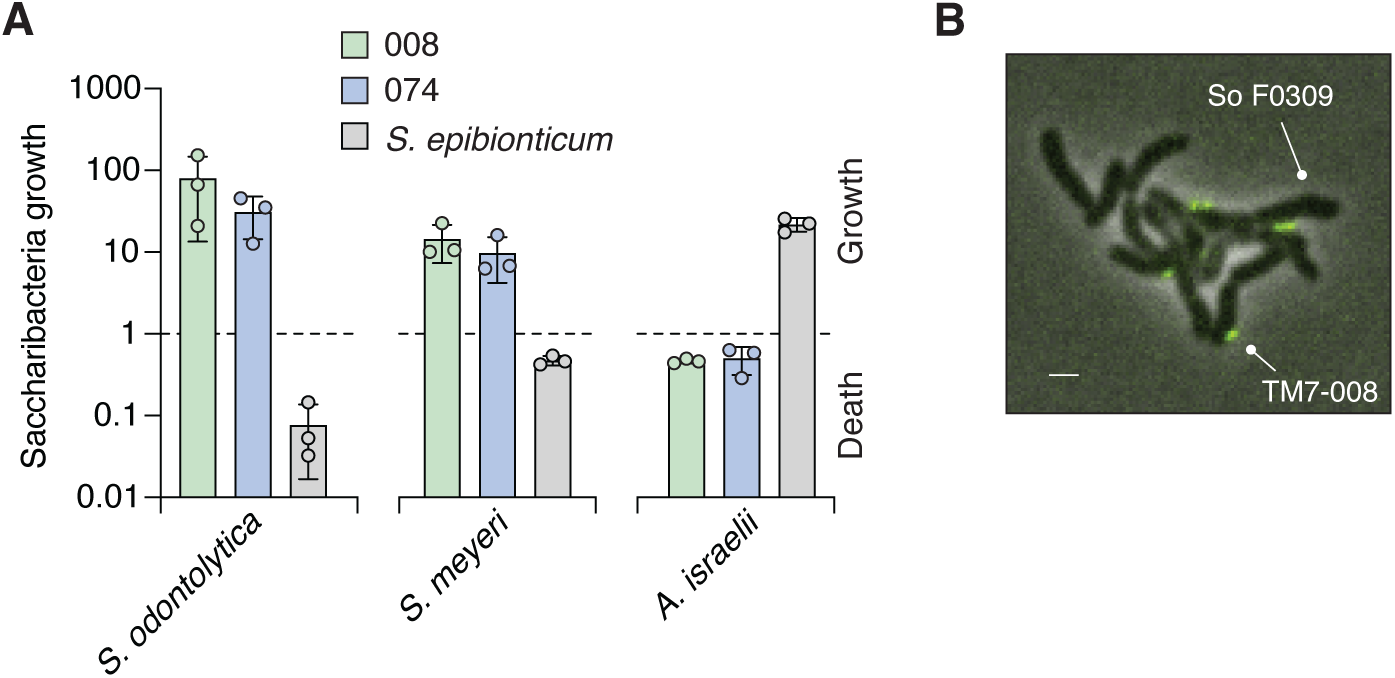
Host species compatibility of Saccharibacterial epibionts. A) Maximum growth or population decline (fold change) observed for TM7-008, TM7-074, and *Southlakia epibionticum* during co-culture with indicated host strains. Data represent mean±SD, n=3 biological replicates. *A. israelii*, *Actinomyces israelii.* B) Fluorescence and phase contrast composite micrograph of a So F0309–TM7-008-*sfgfp* co-culture. Scale bar, 1 µm.

**Supplemental Figure 2.**
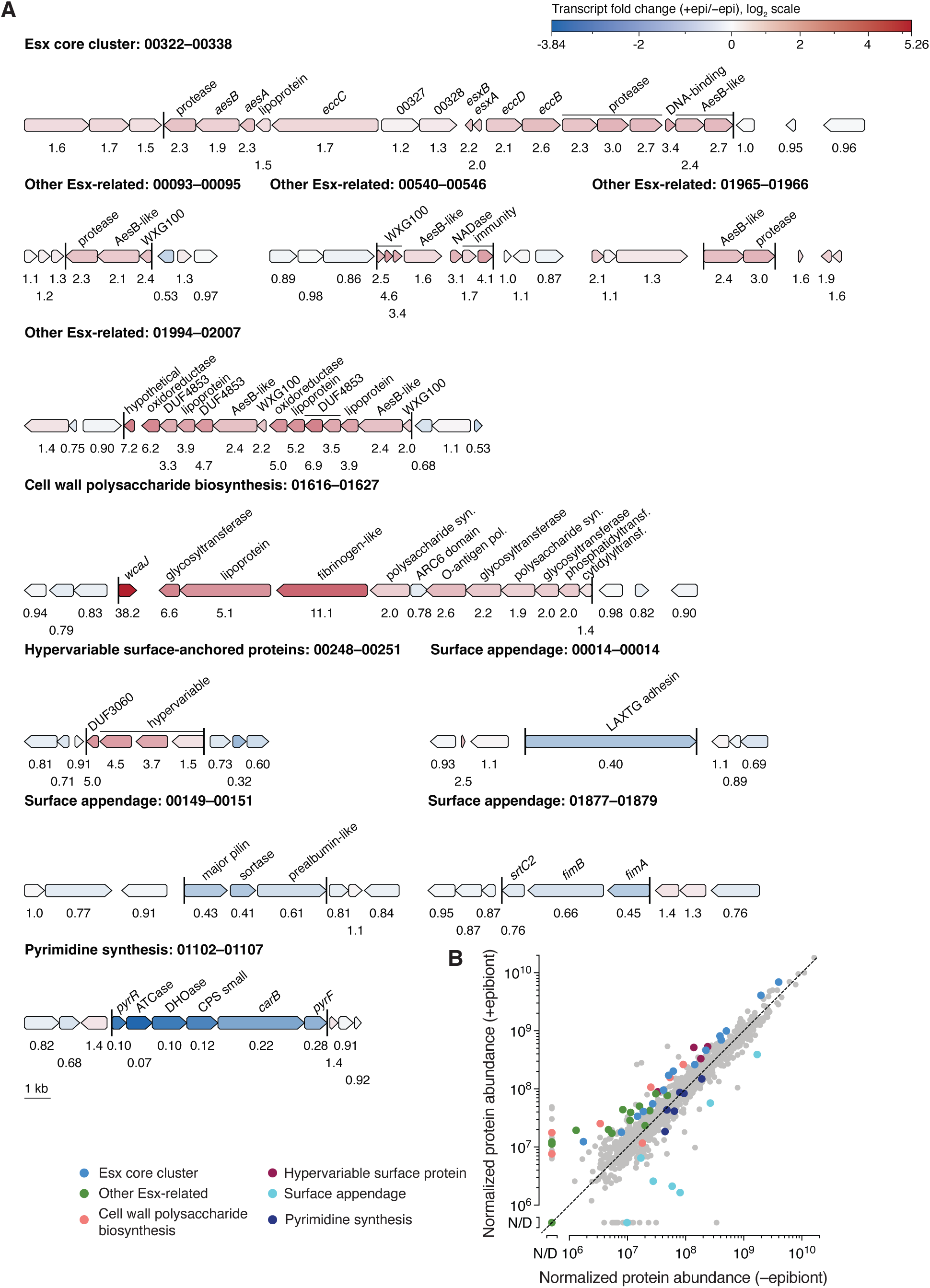
Clusters of genes in *S. odontolytica* F0309 with significantly altered transcription and translation during infection with TM7-008. A) Genes are colored according to their log2-fold change in expression during co-culture with TM7-008 compared to pure culture controls, and flanking genes unaffected by infection with TM7-008 are shown for comparison. Color intensity is scaled to reflect the maximally affected genes. Transcript fold change value is listed below each gene. Locus tags correspond to OBNOAD_xxxxx. B) Scatter plot showing global protein level changes (normalized LFQ intensity) in So F0309 proteins during co-culture with TM7-008 (+epibiont) compared to pure culture controls (-epibiont). LFQ intensity is normalized by summed LFQ intensity of all So F0309 protein groups in each sample. Any normalized LFQ intensity that is <10^6^ is designated as not detected. Proteins are colored in the same way as in Figure 1, and are from the same experiment depicted in Figure 1C. n = 2 biological replicates. N/D, not detected.

**Supplemental Figure 3.**
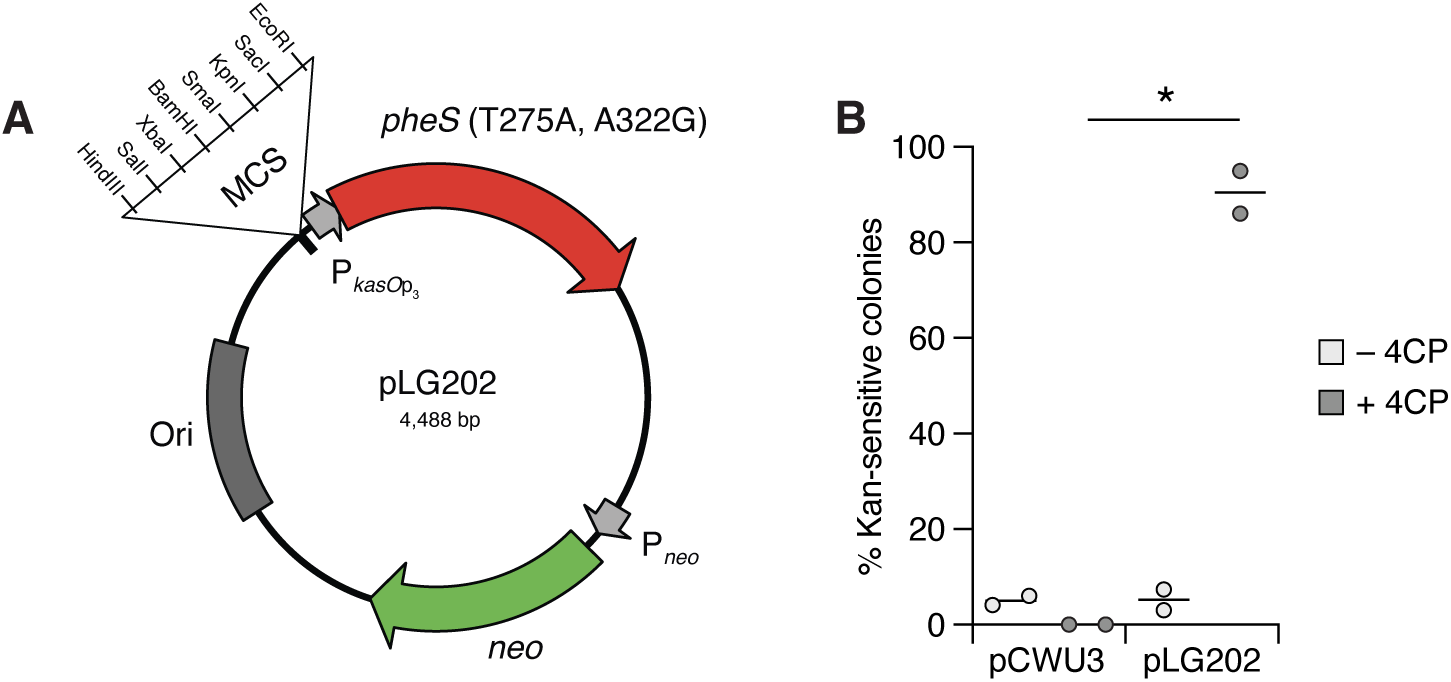
Development of a counterselection-based allelic exchange system for *Schaalia* and related genera. A) Allelic exchange vector map. To generate pLG202, the *mCherry* cassette of pCWU3^57^ was replaced with So F0309 *pheS*^T275A, A322G^ under the control of the strong promoter *kasO*p_3_ ^78^. B) Efficiency of *pheS*-based counterselection. Parental vector pCWU3 and pLG202 containing the same gene fragment for deleting *esrG1* (OBNOAD_00283) were integrated into So F0309 via transformation and plating on selective medium (kanamycin). Merodiploids were grown in non-selective media, then passaged into medium containing 4- chloro-phenylalanine (4CP) to counterselect against the integrated vector backbone. Control cultures were passaged without 4CP. The efficiency of counterselection was assessed by screening resulting clones for kanamycin sensitivity. Data represent the mean, n = 2 biological replicates. Paired two-tailed t test comparing means; *p≤0.05;

**Supplemental Figure 4.**
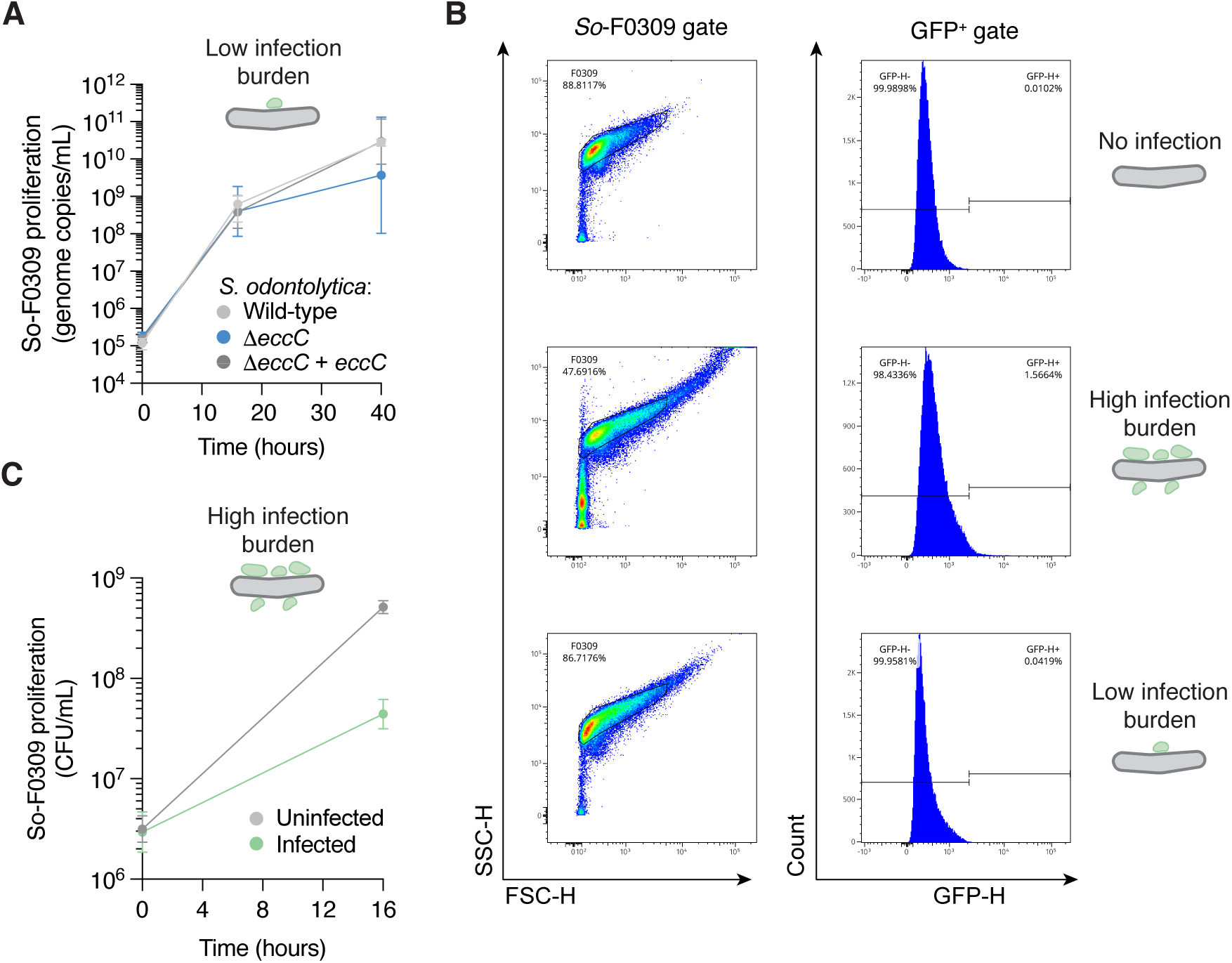
Quantification of *S. odontolytica* F0309 growth and TM7-008 infection levels under high and low infection burden conditions. A) Growth of the indicated So F0309 strains during infection by TM7-008 under a low infection burden condition (see results). Data represent mean±SD, n=3 biological replicates. B) Representative flow cytometry plots detecting the level of TM7-008 infection on So F0309 under high or low infection conditions (see methods). So F0309 events were first identified by forward and side scatter gating, and GFP^+^ events were detected within these gates. A strain of TM7-008 was used that constitutively expresses GFP to enable fluorescent detection of infection burden. C) Growth of So F0309 in the presence or absence of infection by TM7-008 under high burden of infection conditions. Data represent mean±SD, n=3 biological replicates.

**Supplemental Figure 5.**
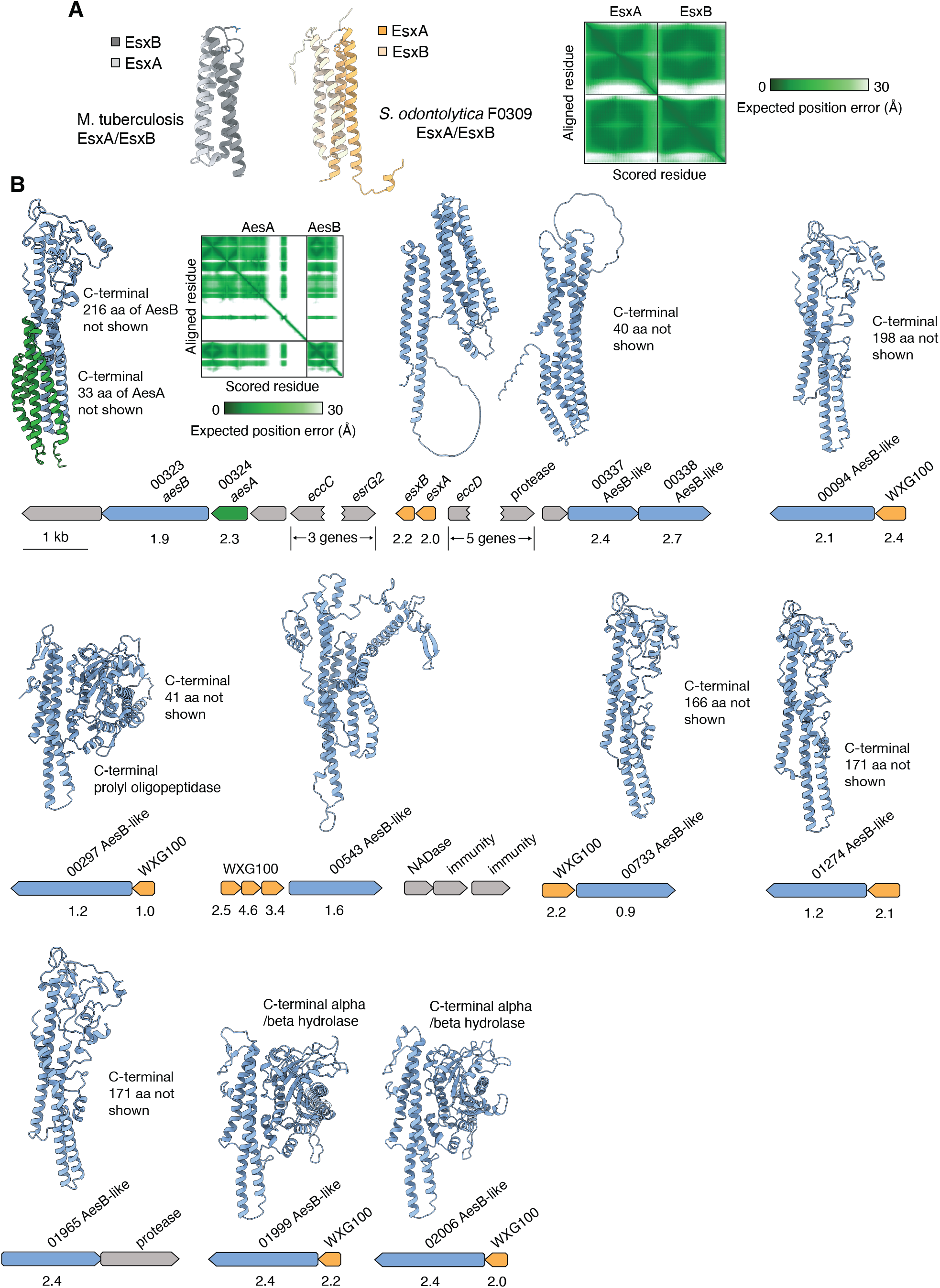
Structural models and genomic arrangement of predicted Esx substrates in So F0309. A) AlphaFold3-predicted model of the complex formed between So F0309 EsxA and EsxB (yellow), compared with a crystal structure of the EsxA/EsxB heterodimer from *Mycobacterium tuberculosis* (grey; PDB 3FAV). Residues forming the conserved WxG secretion motif are shown in stick representation. The predicted aligned error plot for the *S. odontolytica* AlphaFold3 model is shown on the right. B) AlphaFold3 models of predicted Esx substrates in So F0309 and the corresponding genetic loci. The predicted aligned error plot for the AesA–AesB model is included. AesB and AesB-like proteins are colored in blue, AesA in green, and WXG100-like proteins in yellow. Numbers shown below genes represent fold change values for So F0309 transcript abundance in co-cultures with TM7-008 compared to So F0309 mono-cultures.

**Supplemental Figure 6.**
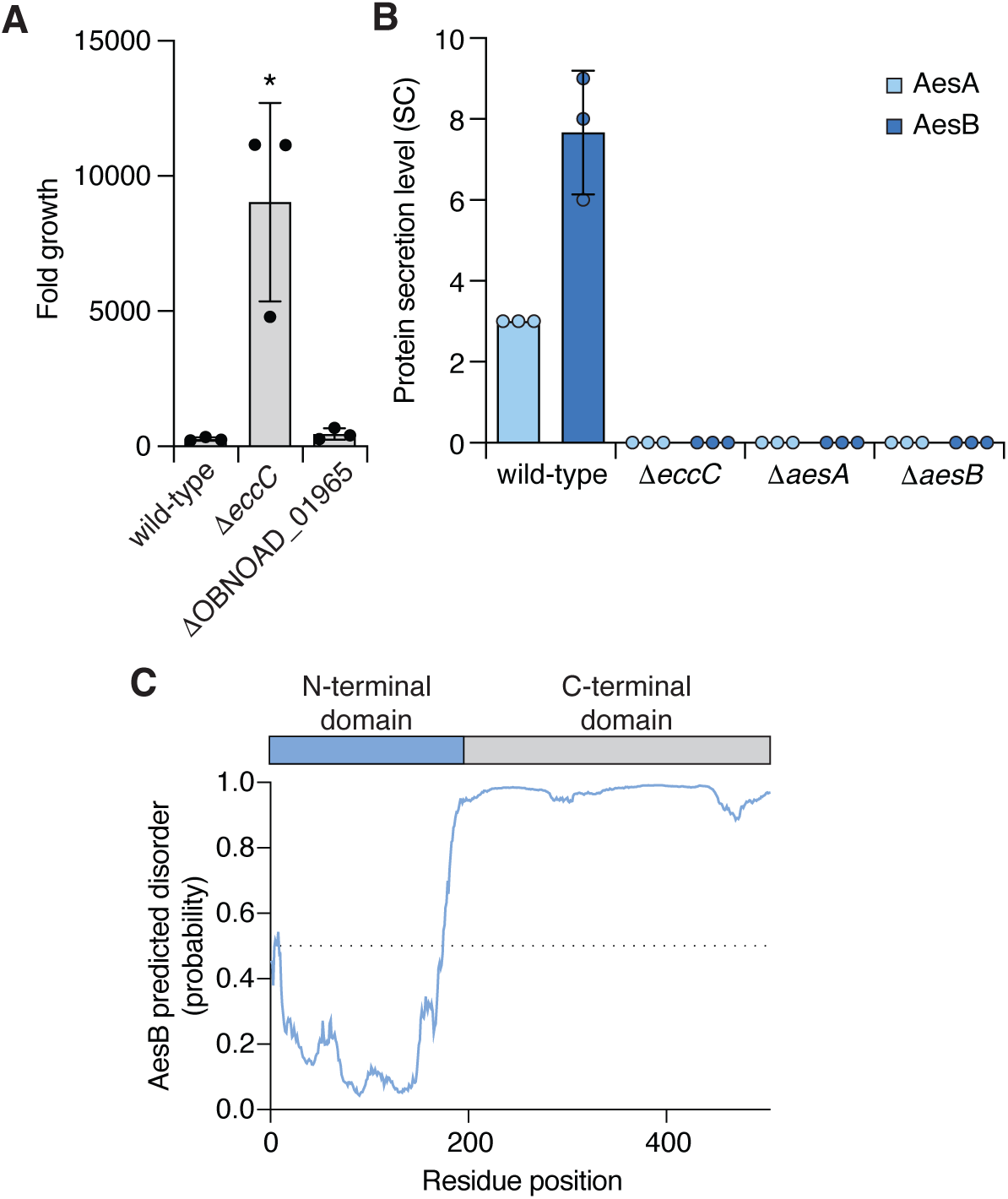
AesA and AesB secretion is mutually dependent and AesB contains a C-terminal IDR. A) Growth of TM7-008 (fold change) achieved on the indicated *S. odontolytica* F0309 strains. OBNOAD_01965 encodes a candidate Esx substrate sharing protein sequence-level homology with AesB. Data represent mean±SD, n=3 biological replicates. *p≤0.05; one-way ANOVA with multiple comparisons to the wild-type mean. B) Level of AesA and AesB detected in cell-free culture supernatant of the indicated So F0309 strains. SC, spectral counts. Data represent the mean ± s.d. (n = 3 technical replicates). C) IUPred3 prediction of disordered regions in AesB^72^. The relative positions of the N- and C-terminal domains are depicted in the schematic above the graph. The dashed line represents a disorder probability of 50%.

**Supplemental Figure 7.**
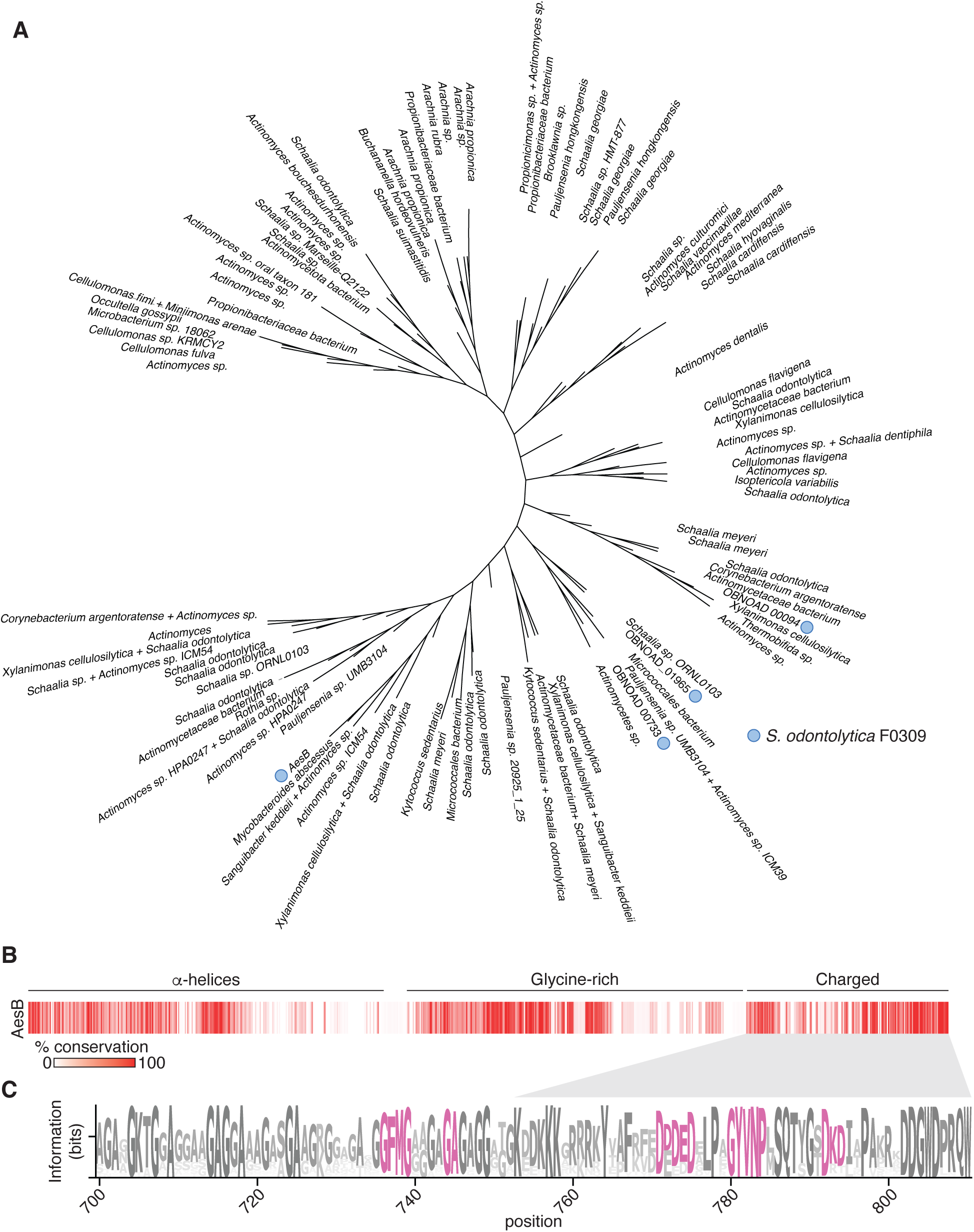
Phylogenetic analysis and sequence diversity of AesB. A) Unrooted maximum likelihood phylogeny of AesB homologs identified by iterative PSI-BLAST searching of the clustered NR database^69^. Proteins found in So F0309 indicated with blue circles. B) Relative conservation at each residue position of AesB, coloured according to the scale at bottom left. Conservation was calculated using the multiple sequence alignment assembled in (A). C) Sequence log deriving from an alignment of the C-terminal region of AesB. The grey wedge indicates the position of the conserved C-terminal charged region. Residues experimentally tested for their importance in inhibiting replication of TM7-008 are highlighted in pink.

**Supplemental Figure 8.**
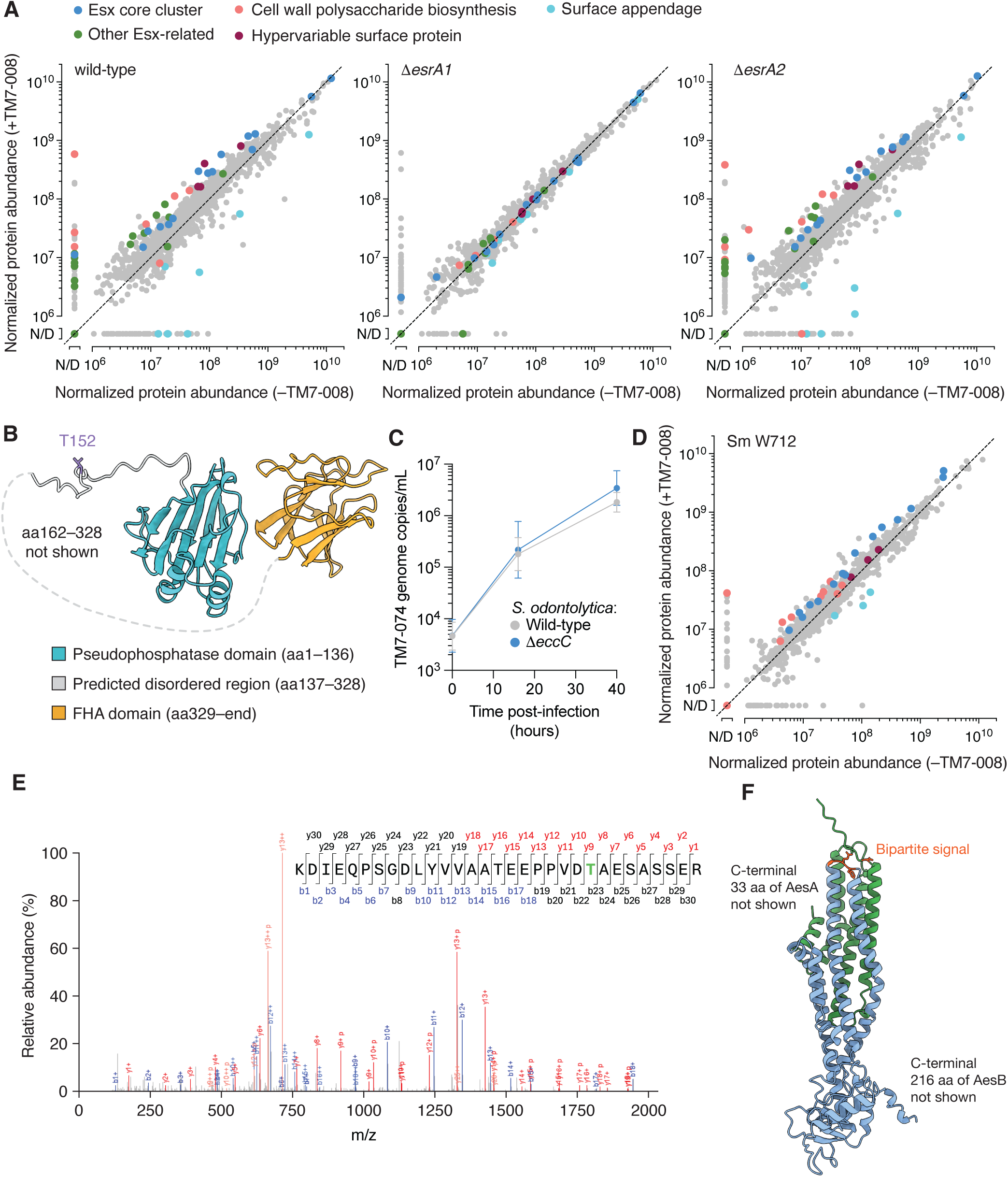
The Esr1-mediated response to TM7-008 infection is conserved in *S. meyeri* W712. A) Scatter plot showing global protein level changes (normalized LFQ intensity) in the indicated strains of So F0309 during co-culture with TM7-008 compared to pure culture conditions. LFQ intensity is normalized by summed LFQ intensity of all So F0309 protein groups in each sample. Any normalized LFQ intensity that is <10^6^ is designated as not detected. Proteins are colored in the same way as in Figure 3D and derive from the same dataset depicted there. n = 2 biological replicates. N/D, not detected. B) AlphaFold3 model of So F0309 EsrG1. The site of phosphorylation (T152) is highlighted in purple. C) Growth of TM7-074 on the indicated So F0309 strains. Data represent mean±SD, n=3 biological replicates. D) Scatter plot showing global protein level changes (normalized LFQ intensity) of *S. meyeri* W712 during co-culture with TM7-008 compared to pure culture. LFQ intensity is normalized by summed LFQ intensity of all *S. meyeri* W712 protein groups in each sample. Any normalized LFQ intensity that is <10^6^ is designated as not detected. Proteins are colored in the same way as in Figure 4A. n = 3 technical replicates. E) Tandem mass spectra of the indicated *S. meyeri* W712 peptide derived from co-culture with TM7-008. Matched fragmentation ions (N-terminal b-ions in blue and C-terminal y-ions in red) and the site of phosphorylation (green; T263 in EsrG1) are indicated. F) AlphaFold3 model of the AesA–AesB heterodimer (AesA green; AesB blue) with the bipartite signal motif highlighted in orange red.

